# VEGF and FGF signaling during head regeneration in hydra

**DOI:** 10.1101/596734

**Authors:** Anuprita Turwankar, Surendra Ghaskadbi

**Affiliations:** Developmental Biology Group, MACS-Agharkar Research Institute, Savitribai Phule Pune University, G.G. Agarkar road, Pune 411004, India

**Keywords:** hydra, decapitation, head regeneration, FGF-1, FGFR-1, VEGFR-2

## Abstract

**Background:** Vascular endothelial growth factor (VEGF) and fibroblast growth factor (FGF) signaling pathways play important roles in the formation of the blood vascular system and nervous system across animal phyla. We have earlier reported VEGF and FGF from *Hydra vulgaris* Ind-Pune, a cnidarian with a defined body axis, an organized nervous system and a remarkable ability of regeneration. We have now identified three more components of VEGF and FGF signaling pathways from hydra. These include *FGF-1*, FGF receptor 1 (*FGFR-1*) and VEGF receptor 2 (*VEGFR-2*) with a view to deciphering their possible roles in regeneration.

**Methods:** *In silico* analysis of proteins was performed using Clustal omega, Swiss model, MEGA 7.0, etc. Gene expression was studied by whole mount in situ hybridization. VEGF and FGF signaling was inhibited using specific pharmacological inhibitors and their effects on head regeneration were studied.

**Results:** Expression patterns of the genes indicate a possible interaction between FGF-1 and FGFR-1 and also VEGF and VEGFR-2. Upon treatment of decapitated hydra with pharmacological inhibitor of FGFR-1 or VEGFR-2 for 48 hours, head regeneration was delayed in treated as compared to untreated, control regenerates. When we studied the expression of head specific genes *HyBra1* and *HyKs1* and tentacle specific gene *HyAlx* in control and treated regenerates using whole mount in situ hybridization, expression of all the three genes was found to be adversely affected in treated regenerates.

**Conclusions:** The results suggest that VEGF and FGF signaling play important roles in regeneration of hypostome and tentacles in hydra.

## 1. Introduction

The phenomenon of regeneration has intrigued biologists since long. Regeneration is a process of tissue replacement and animals can regenerate either through epimorphosis which involves active cellular proliferation and blastema formation or morphallaxis which occurs through remodeling of the existing tissue [1]. Regeneration essentially involves wound healing, dedifferentiation (morphallaxis) or dedifferentiation followed by proliferation of the cells (epimorphosis) that leads to replacement of the lost structure(s) [2]. Animal regeneration studies date back to the 16th century when René-Antoine Ferchault de Réaumur, a French entomologist presented his detailed study of crayfish claw regeneration in the French Academy. In 1744, Abraham Trembley, a Swiss naturalist published his studies on hydra regeneration, budding and tissue grafting. He demonstrated that small pieces of hydra polyp gave rise to the complete polyp. Lazzaro Spallanzani, in 1768, described limb and tail regeneration in newts and tadpoles [3]. The regenerative abilities vary greatly in the animal kingdom; organisms belonging to the basal phyla like cnidarians (e.g. hydra) and platyhelminths (e.g. planaria) are capable of whole body regeneration from a small body fragment. On the other hand, zebrafish can regenerate fins and heart whereas newts are capable of limb and lens regeneration. Mammals have relatively very limited ability to regenerate and exhibit regeneration of injured tissues like liver, pancreas and heart [4]. In contrast, other organisms like birds, nematodes and leeches are hardly capable of any regeneration [5].

Hydra, a freshwater polyp and a member of phylum Cnidaria, has been a popular model organism to study pattern formation because of its remarkable regeneration capability. Hydra is a diploblastic animal, in which ectoderm and endoderm are separated by mesoglea, made up of extracellular matrix. Hydra exhibits radial symmetry with a distinct oral-aboral axis and represents one of the first animals with a defined body axis and a simple nervous system in the form of a nerve net [6]. Hydra polyp resembles a perpetual embryo that does not exhibit organismal senescence. This lack of senescence in hydra is attributed to its multipotent stem cells. Three stem cell lineages are present in hydra; these include ectodermal and endodermal epithelial stem cells that give rise to the respective epithelial cells and interstitial stem cells (i cells) that give rise to the gland cells, nematocytes, nerve cells and gametes. The i cells reside in the upper two third of body column of hydra. Interstitial cells continuously proliferate and give rise to differentiated cells, which migrate towards the tentacles and basal disc of the polyp. Thus, terminally differentiated cells are present at the two extremities of hydra [7]. Hydra has a tremendous potential for regeneration. If a hydra is cut in several pieces, each piece, except for the two extremities, will regenerate and give rise to a new polyp. Also, the original polarity of hydra is maintained in the regenerated polyp; hence regeneration is tightly regulated in hydra [8]. Head regeneration is brought about by two different mechanisms depending on the position of the injury. When amputated at about 50% body length (mid-gastric bisection), hydra undergoes basal head regeneration while if amputated at about 80% body length (decapitation), it undergoes apical head regeneration. In case of mid-gastric bisection, cells of the interstitial lineage surrounding the cut site undergo apoptosis and these apoptotic cells are engulfed by the endodermal epithelial cells. The apoptotic cells act as a source of Wnt3 and activate the Wnt-β-catenin pathway. This mode of regeneration resembles epimorphosis. In decapitated hydra, the epithelial cells at the cut site upregulate Wnt3 expression that leads to remodelling of the pre-existing tissue to regenerate the lost head, in the absence of cell proliferation. Thus, apical head regeneration exhibits morphallactic mode of regeneration [2].

Hydra regeneration requires cell-cell communication effected through various cell signaling pathways. These pathways are conserved as similar pathways are functional in the vertebrate embryo [9]. One such conserved cell signaling pathway includes Receptor tyrosine kinases (RTKs) [10, 11]. The ligands employed by these recep tors are usually a number of growth factors. FGF and VEGF are two such growth factors that signal through the RTKs - FGF receptor (FGFR) and VEGF receptor (VEGFR), respectively. FGFs play a role in regeneration of limb (frog), tail (axolotl), fin (zebrafish), lens (newt), retina (chick and zebrafish), components of the nervous system (zebrafish), skeletal muscles (mouse), and bone (mouse) in vertebrates. FGF signaling is also involved in lung, intestine and liver regeneration of vertebrates [12]. Role of VEGF signaling in regeneration of liver, lungs and bone marrow vessels has been demonstrated [13]. A FGFR like gene – kringelchen that belongs to the RTK family, is involved in boundary formation and bud detachment in hydra [14]. Another FGF identified in hydra – FGFf, is thought to play a role in cell movement and morphogenesis [15].

Homologues of vascular endothelial growth factor (VEGF) and fibroblast growth factor (FGF) in hydra have been reported from our laboratory [16]. These two angiogenic factors are of particular importance and interest in hydra, because hydra is a simple metazoan in which the mesoderm (and hence angiogenesis) is completely absent. Presence of VEGF and FGF in hydra therefore suggests these molecules are important for processes other than angiogenesis. As a part of our continues efforts to understand roles of VEGF and FGF signaling in hydra, particularly in regeneration, we have identified, isolated and partially characterized HyFGF-1, HyFGFR-1 and HyVEGFR-2 from *Hydra vulgaris* Ind-Pune. *In silico* analysis and sequence comparison of these genes with their vertebrate counterparts reveal that these genes are conserved during evolution. Since the spatiotemporal expression pattern of a gene often provides insights into its function in morphogenesis, localization of these genes in hydra was studied using whole mount in situ hybridization. The expression patterns indicate interaction between the respective ligands and receptors (HyFGF-1 and HyFGFR-1; HyVEGF and HyVEGFR-2). Inhibition of the receptor tyrosine kinase activity of HyFGFR-1 and HyVEGFR-2 with specific pharmacological inhibitors resulted in delayed head regeneration which was evident from morphology as well as expression of head specific marker genes in regenerating polyps. The present results thus suggest role of VEGF and FGF signaling during apical head regeneration in hydra.

## 2. Materials and Methods

### 2.1. Hydra culture and maintenance

Clonal cultures of *Hydra vulgaris* Ind-Pune [11] were maintained in glass crystallizing dishes containing hydra medium, composed of 1mM CaCl_2_, 0.1mM MgSO_4_, 0.1mM KCl, 1mM NaCl and 1mM Tris Cl, pH 8. Hydra were maintained at constant temperature of 18°C with 12 hr light/dark cycle. The polyps were fed on alternate days with *Artemia salina* nauplii. Hatching of *Artemia* cysts was done in artificial sea water. Freshly hatched larvae were collected, washed thoroughly with water and used for feeding. 7 hours post feeding, hydra were washed thoroughly and old medium was replaced with fresh hydra medium.

### 2.2. Study of morphology of regenerating hydra

Groups of 10 non-budding, 24 hrs starved hydra were decapitated and allowed to regenerate for different time periods - 0.5, 1.5, 3, 6, 12, 24, 48 and 72h. Polyps were observed under bright field illumination with Olympus SZX16 stereo microscope and photographed using an Olympus DP71 camera.

### 2.3. Isolation and cloning of *FGF-1, FGFR-1* (partial and complete CDS) and *VEGFR-2* (partial and complete CDS) homologues from *Hydra vulgaris* Ind-Pune

Total RNA was extracted from hydra using TriReagent (Sigma, USA) and cDNA was synthesized using Verso cDNA synthesis kit (Thermo Fisher Scientific, USA). Predicted gene sequence of *FGF-1* from *Hydra magnipapillata* (GenBank accession no. XM_004209493.2) and predicted gene sequences of *FGFR-1* (GenBank accession no. NM_001309675.1) and *VEGFR-2* (GenBank accession no. XM_012699367.1) from *Hydra vulgaris* were retrieved from NCBI database. These sequences were used as a template to design oligonucleotide primers for amplification and isolation of the respective sequences from *Hydra vulgaris* Ind-Pune. The details of the primer sequences are as follows:

*FGF-1*:

FW: 5’ ATG ATA TTG CTT CAA AGT TTT TTT GAG 3’

REV: 5’ TTA TGC TTT CTG CTT TTT TCC ACC 3’

*FGFR-1* partial CDS:

FW: 5’ CCA AAA AGT TCT GAA GTG ATT GC 3’

REV: 5’ GGG ATC ACC TTC ATC AAT TAT ACG 3’

*VEGFR-2* partial CDS:

FW: 5’ GCC ATT GTC GCT TCA CTT GG 3’

REV: 5’ TTT GCA TGC GGA ACG AGA AC 3’

*FGFR-1* complete CDS:

FW: 5’ ATG ATG TTG TTT TTG TGT TTG GTT TC 3’

REV: 5’ TTA AAC CGG CAA ATT GTC AAA AGG 3’

*VEGFR-2* complete CDS:

FW: 5’ ATG TTA CGA TAC TTT CTA GTT TTA ATT TAC TGG 3’

REV: 5’ TTA ATT CAA GTT TGC GTA CAT AGT AGT AGC AC 3’

PCR conditions used for amplification were as follows: Initial denaturation at 94°C for 4 min followed by 40 cycles of denaturation at 94°C for 30 sec, annealing for 40 sec at 56.7°C for *FGF-1* and at 60°C for both *FGFR-1* and *VEGFR-2* partial CDS and extension at 72°C for 50 sec and final extension at 72°C for 5 min. The complete CDS of *FGFR-1* and *VEGFR-2* were amplified using Platinum® *Taq* DNA Polymerase High Fidelity (Thermo Fisher Scientific, USA). PCR conditions used for amplification were as follows: Initial denaturation at 95°C for 2 min followed by 40 cycles of denaturation at 94°C for 10 sec, annealing for 1 min at 63.8°C and extension at 68°C for 2 min 30 sec for *FGFR-1* and for 4 min 30 sec for *VEGFR-2*. Final extension was carried out at 68°C for 10 min. The amplified PCR products were purified and cloned into pGEM-T Easy (Promega, Madison, WI, USA) and sequenced (1^st^ Base, Malaysia).

### 2.4. *In silico* analysis of HyFGF-1, HyFGFR-1 and HyVEGFR-2

Putative protein sequences of HyFGF-1, HyFGFR-1 and HyVEGFR-2 were analyzed by SMART to determine the functional domains [36]. EMBL-EBI Clustal omega software was used for multiple sequence alignment of the protein sequences in order to determine their identity with the known proteins [37]. Other important amino acid residues within the protein were assigned manually based on available literature. Homology based putative protein models were generated for HyFGF-1, HyFGFR-1 and HyVEGFR-2 using Swiss Model program at ExPaSy Server [38]. These models were compared with the solved structures from other organisms to determine homology using Swiss PDB Viewer Software Deep View [39]. The models were superimposed using ‘Iterative magic fit’ and the degree of similarity was determined using the Root Mean Square Deviation (RMSD) values. Phylogenetic trees were constructed by Neighbor Joining method using MEGA7 [40]. Bootstrap analysis with 5000 replicates was carried out. The protein sequences used for constructing the phylogenetic tree were obtained from NCBI and UniProt. FGF-1 sequences obtained from NCBI are - *Amphimedon queenslandica* (XP_003387084.1), *Hydra vulgaris* Ind-Pune (AND74488.1), *Nematostella vectensis* (ABN70833.1), *Tribolium castaneum* (XP_008196985.1), *Danio rerio* (AAH59588.1), *Alligator sinensis* (XP_006023603.1), *Chelonia mydas* (XP_007063672.1) and *Gallus gallus* (XP_015149497.1). *Xenopus laevis* (Q6GLR6), *Mus musculus* (P61148) and *Homo sapiens* (P05230) were taken from UniProt.

Sequences of FGFR-1 obtained from NCBI are - *Amphimedon queenslandica* (XP_019849350.1), *Hydra vulgaris* Ind-Pune (AZQ04902.1), *Nematostella vectensis* (ABO92763.1) and *Chelonia mydas* (XP_027680214.1). *Ciona intestinalis* (Q4H3K6), *Drosophila melanogaster* (Q07407), *Danio rerio* (Q90Z00), *Xenopus laevis* (P22182), *Alligator sinensis* (A0A1U7RGJ6), *Gallus gallus* (P21804), *Mus musculus* (P16092) and *Homo sapiens* (P11362) were taken from UniProt.

Sequences of VEGFR-2 obtained from NCBI are - *Amphimedon queenslandica* (XP_011406831.2), *Hydra vulgaris* Ind-Pune (AZQ04903.2), Idiosepius paradoxus (BAI67804.1), *Xenopus laevis* (XP_018085457.1) and *Gallus gallus* (AAR26285.1). Sequences of *Podocoryna carnea* (Q674V1), *Drosophila melanogaster* (Q8IPG1), *Danio rerio* (Q5GIT4), *Chelonia mydas* (M7AKY2), *Alligator sinensis* (A0A1U7RRK7), *Mus musculus* (P35918) and *Homo sapiens* (P35968) were taken from UniProt.

### 2.5. Whole mount *in situ* hybridization

Whole mount *in situ* hybridization was carried out to localize transcripts of *FGF-1, FGFR-1* and *VEGFR-2* as described by Krishnapati and Ghaskadbi (2013), with a few modifications. Briefly, the sense and antisense riboprobes for *FGF-1, FGFR-1* and *VEGFR-2* were synthesized by *in vitro* transcription reaction (Roche). Hydra polyps were starved for 48 hrs, relaxed in 2% urethane for 1-2 min and fixed in 4% paraformaldehyde overnight at 4°C. The polyps were washed with ethanol until the polyps lost colour. The polyps were rehydrated by subsequent washes with 75%, 50% and 25% ethanol in 1X PBST (Phosphate buffered saline Tween). Polyps were permeabilized with 10 μg/ml proteinase K at room temperature for 10 min to facilitate the entry of probe into the cells. Proteinase K activity was stopped by addition of 1X Glycine in PBST for 10 min. Further, Glycine in PBST was exchanged with triethanolamine for 10 min to reduce background staining, followed by triethanolamine+acetic acid wash for 10 min. PBST washes were carried out and hydra were refixed in 4% paraformaldehyde at 4°C overnight. The polyps were washed with PBST and 2X SSC successively and equilibrated in prehybridization buffer for 10 min. The polyps were transferred to fresh prehybridization buffer to block nonspecific sites at 60°C for 3 hrs. Hybridization was carried out by the addition of appropriate DIG labelled sense and antisense riboprobes at the same temperature for 2.5 days. The polyps were washed with hybridization buffer for 10 min followed by graded series of hybridization buffer and 2X SSC as follows:

100% Hybridization solution, 75% Hybridization solution + 25% 2X SSC, 50% Hybridization solution + 50% 2X SSC, 25% Hybridization solution + 75% 2X SSC and 100% 2X SSC; each for 10 min. Stringency washes of 0.5X SSC and 0.1% CHAPS solution was carried out at 60°C (3 washes of 20 min each) to remove any unbound and non-specifically bound probe. MABT washes were given for about an hour and non-specific protein binding sites were blocked by incubating the polyps in 20% fetal bovine serum (FBS) in MABT at room temperature for 3 hrs. This was followed by overnight incubation in anti-DIG antibody (1:3000) at 4°C. Excess antibody was removed by repeated washes of MABT at room temperature (8 washes of 20 min each). The polyps were incubated in MABT at room temperature on a rocker incubator overnight. The polyps were equilibrated in NTMT (pH-9.5) followed by NTMT-Levamisole for 5 min to reduce background alkaline phosphatase activity. Fresh NTMT was added with NBT and BCIP solution and incubated in dark. After development of colour, the polyps were transferred to methanol and stored at −20°C. Stained polyps were observed under incident light with Olympus SZX16 stereo microscope and photographed using an Olympus DP71 camera.

### 2.6. Treatment with SU5402 and SU5416

SU5402 is a fibroblast growth factor receptor 1 (FGFR1) inhibitor [34] while SU5416 is a potent and specific inhibitor of VEGF receptor tyrosine kinase 2 (Flk-1/KDR) [35]. Whole, non-budding hydra were treated with different doses of inhibitors to determine effective doses resulting in maximum abnormality and minimum mortality. Based on this criteria, 20 µM of SU5416 and 40 µM of SU5402 (Calbiochem) were chosen for treatment of regenerating hydra. Hydra were decapitated and allowed to regenerate head for 48 hrs in the presence or absence of SU5416 and SU5402. Inhibitor solution was replaced with fresh inhibitor solution after 24 hrs. Regenerating hydra kept in hydra medium served as master controls while those kept in DMSO solution served as solvent controls. After treatment for 48 hrs, the polyps were transferred to fresh hydra medium for recovery for a further 48 hrs. After recovery, polyps were observed and photographed as before. For whole mount *in situ* hybridization post inhibitor treatment, polyps were fixed overnight and whole mount *in situ* hybridization was carried out as before.

### 2.7. Identification, isolation, cloning and expression of head and tentacle marker genes from *Hydra vulgaris* Ind-Pune

To monitor head regeneration after inhibitor treatment, expression patterns of head specific genes – *HyBra1, HyKs1* and tentacle specific gene - *HyAlx* were studied. Total RNA was extracted from hydra using TriReagent (Sigma, USA) and cDNA was synthesized using Verso cDNA synthesis kit (Thermo Fisher Scientific, USA). The nucleotide sequences of *HyBra1* (GenBank accession number: AY366371.1), *HyKs1* (GenBank accession number: X78596.1) and *HyAlx* (GenBank accession number: AF295531.1) from *Hydra vulgaris* were retrieved from NCBI database. These sequences were used as templates to design oligonucleotide primers for amplification and isolation of the respective sequences from *Hydra vulgaris* Ind-Pune. The details of the primer sequences are as follows:

*HyBra1*:

FW: 5’ ATG AAT GCA AAA GAC ATT GAT GG 3’

REV: 5’ TTA TAT ATT GGA GGG ATA AAC TAG AG 3’

*HyKs1*

FW: 5’ ATG AAA CTA ATA ATT GTG CTT GTA ATG 3’

REV: 5’ TTA AAA ATT CAG GTT GAA TTT TTT TTT AAA G 3’

*HyAlx*

FW: 5’ ATG ATA CAC AAA CCT ATG GC 3’

REV: 5’ TTA ATG AAA ATA ACT ATA TCT TAA AG 3’

PCR conditions used for amplification were: Initial denaturation at 94°C for 4 min followed by 40 cycles of denaturation at 94°C for 30 sec, annealing for 40 sec at 60°C and extension at 72°C for 50 sec and final extension at 72°C for 5 min. The amplified PCR products were purified and cloned into pGEM-T Easy (Promega, Madison, WI, USA) and sequenced (1^st^ Base, Malaysia). Transcripts of *HyBra1, HyKs1* and *HyAlx* in adult, non-budding hydra, were localized by whole mount *in situ* hybridization as before.

## 3. Results

### 3.1. Head regeneration in hydra

Within 0.5 to 1 hour post decapitation (hpd), the two epithelial layers of hydra fused together at the cut site and sealed the apical end. Unlike the columnar cells in the body column of hydra, the cells at the cut site appeared flattened due to the absence of extracellular matrix. At 24-48 hpd, tentacles began to emerge and a fully functional head regenerated after 72 hpd (Fig.1).

**Fig. 1.**
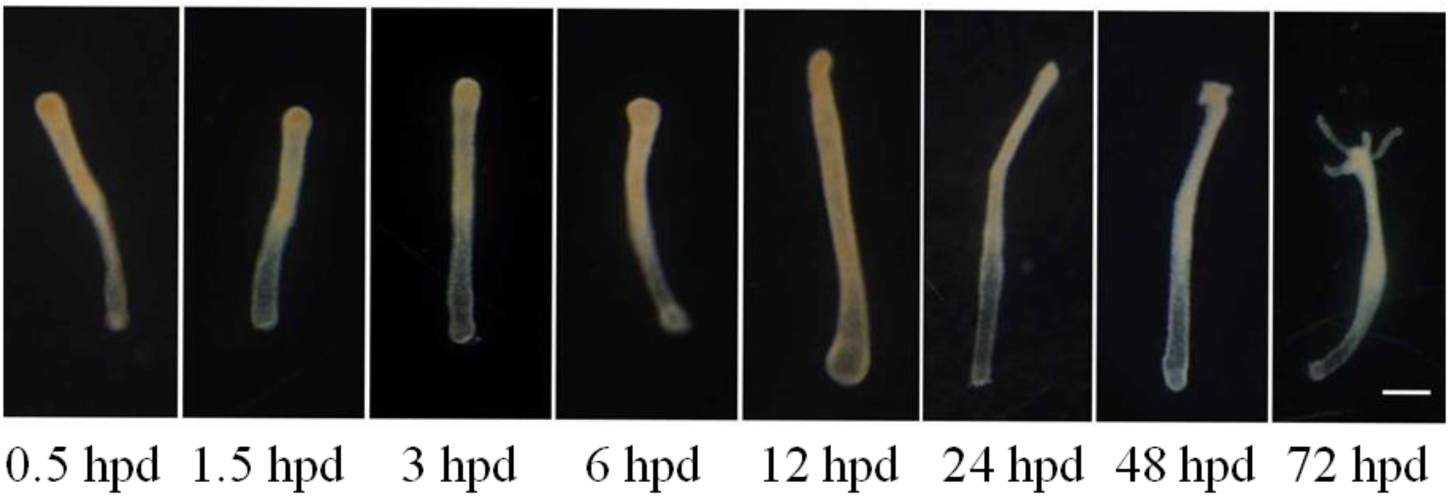
Morphology of hydra during head regeneration, post decapitation. Hydra were decapitated and kept in hydra medium for head regeneration. Morphology of regenerating hydra was noted at different hours post decapitation (hpd). Scale bar, 200 µm.

### 3.2. Isolation of *FGF-1, FGFR-1* and *VEGFR-2* homologues from *Hydra vulgaris* Ind-Pune

*FGF-1* (555 bp), *FGFR-1* (partial CDS – 1020 bp and complete CDS – 2259 bp) and *VEGFR-2* (partial CDS – 905 bp and complete CDS – 4617 bp) were cloned from *Hydra vulgaris* Ind-Pune. Total RNA was extracted and used as a template for cDNA synthesis, followed by gene specific PCR. The clones were sequenced and submitted to NCBI database (https://www.ncbi.nlm.nih.gov/) with GenBank accession numbers as follows: *FGF-1* complete CDS: KU248484; *FGFR-1* partial CDS: MF138882; *VEGFR-2* partial CDS: MF138881; *FGFR-* 1 complete CDS: MH194568 and *VEGFR-2* complete CDS: MH194569. Consistent with the fact that hydra genome is AT rich, A+T content of *FGF-1, FGFR-1 a*nd *VEGFR-2* was found to be 68.64%, 67.1% and 66.1%, respectively.

### 3.3. Functional domains of HyFGF-1, HyFGFR-1 and HyVEGFR-2 proteins are conserved across phyla

Analysis of HyFGF-1, HyFGFR-1 and HyVEGFR-2 proteins with Simple Modular Architecture Research Tool (SMART) revealed the presence of extracellular immunoglobulin domains, transmembrane domains and functional domains within the proteins. HyFGF-1 showed 124 amino acid long FGF domain (Fig. 2a, b), HyFGFR-1 showed the presence of three extracellular immunoglobulin domains, a transmembrane domain and 276 amino acids long tyrosine kinase domain (Fig. 4a, b) while HyVEGFR-2 showed eight extracellular immunoglobulin domains, a transmembrane domain and 317 amino acids long tyrosine kinase domain (Fig. 6a, b). Multiple sequence alignment (MSA) by Clustal Omega analysis showed that the functional FGF domain of HyFGF-1 (Fig. 2c), tyrosine kinase domains of HyFGFR-1 (Fig. 4c) and HyVEGFR-2 (Fig. 6c) were conserved across phyla from hydra to human. HyFGF-1 showed maximum identity of 28.86 % with FGF-1 from chick, followed by 28.19% identity with human FGF-1. The Tyrosine kinase (TyrKc) domain of HyFGFR-1 and human FGFR-1 exhibited maximum identity of 40.73%. TyrKc domain of HyVEGFR-2 and *Podocoryna carnea* VEGFR showed 52.1% identity.

**Fig. 2.**
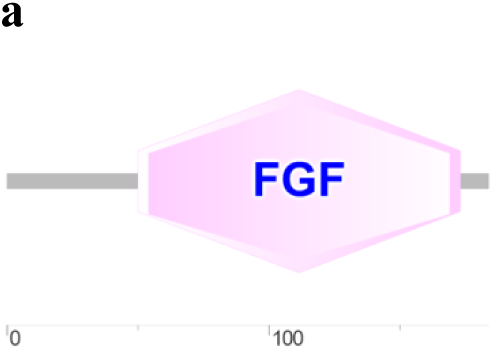
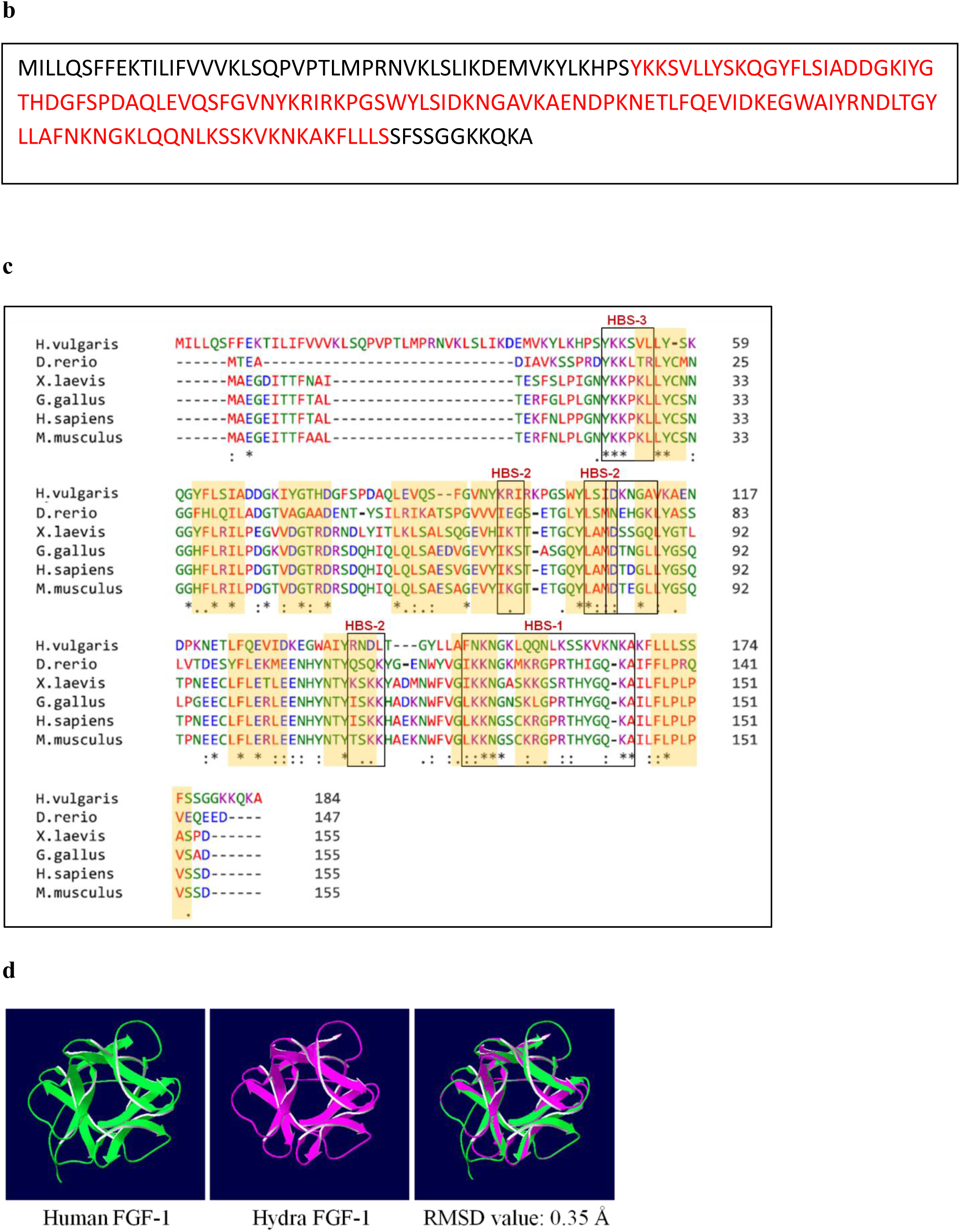
SMART analysis, FGF-1 domain, multiple sequence alignment and homology modelling of FGF-1. (a) SMART analysis of FGF-1 protein shows the characteristic FGF-1 domain. (b) The amino acid sequence of FGF-1 domain is highlighted in red. (c) Multiple sequence alignment of hydra FGF-1 protein with other FGF-1 homologues shows the conservation of amino acid residues – (*) indicates positions which have a single, fully conserved residue, (:) indicates conservation between groups of strongly similar properties and (.) indicates conservation between groups of weakly similar properties. The boxes denote the heparan sulphate glycosaminoglycan (HSGAG) binding sites (HBS) within FGF-1 proteins. The highlighted portions in yellow represent the 12 β strands of the β-trefoil structure. (d) Tertiary structure of HyFGF-1 was simulated from available solved structure of human FGF-1, using Swiss Model Tool. These structures were superimposed using Iterative Magic Fit tool in SPDBV (RMSD: 0.35 Å).

**Fig. 3.**
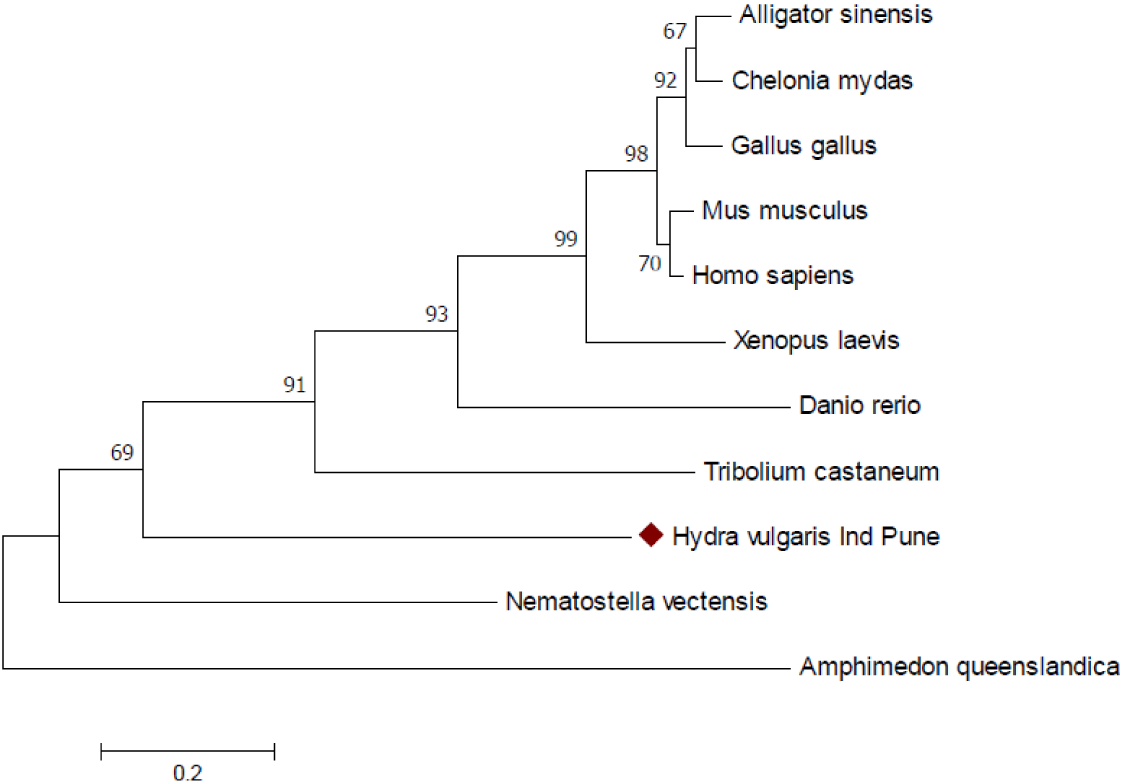
Phylogenetic analysis of FGF-1 conducted in MEGA7. The evolutionary history was inferred using the Neighbor-Joining method. The optimal tree with the sum of branch length = 3.92664366 is shown. The percentage of replicate trees in which the associated taxa clustered together in the bootstrap test (5000 replicates) are shown next to the branches. The tree is drawn to scale, with branch lengths in the same units as those of the evolutionary distances used to infer the phylogenetic tree. The evolutionary distances were computed using the Poisson correction method and are in the units of the number of amino acid substitutions per site. The analysis involved 11 amino acid sequences. All positions containing gaps and missing data were eliminated. There were a total of 101 positions in the final dataset.

**Fig. 4.**
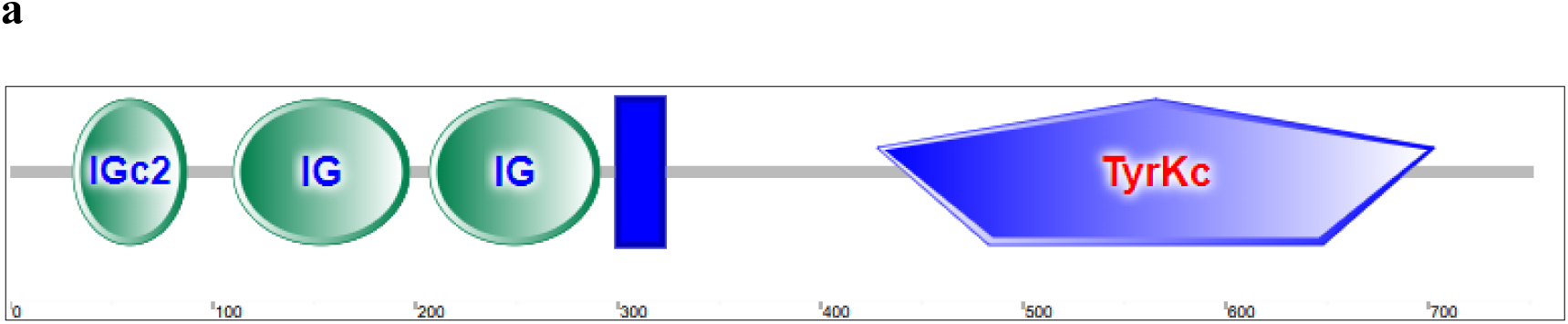
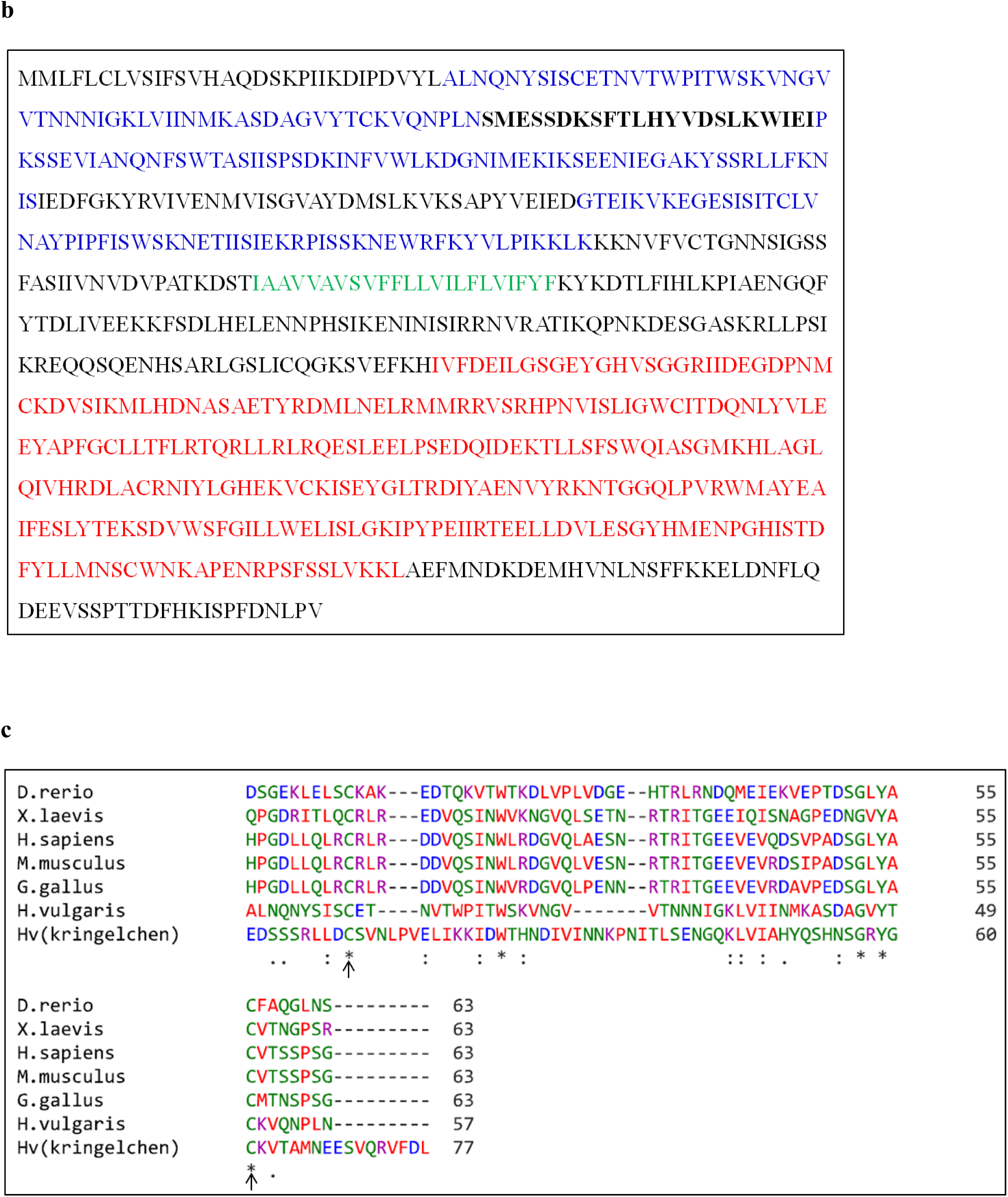
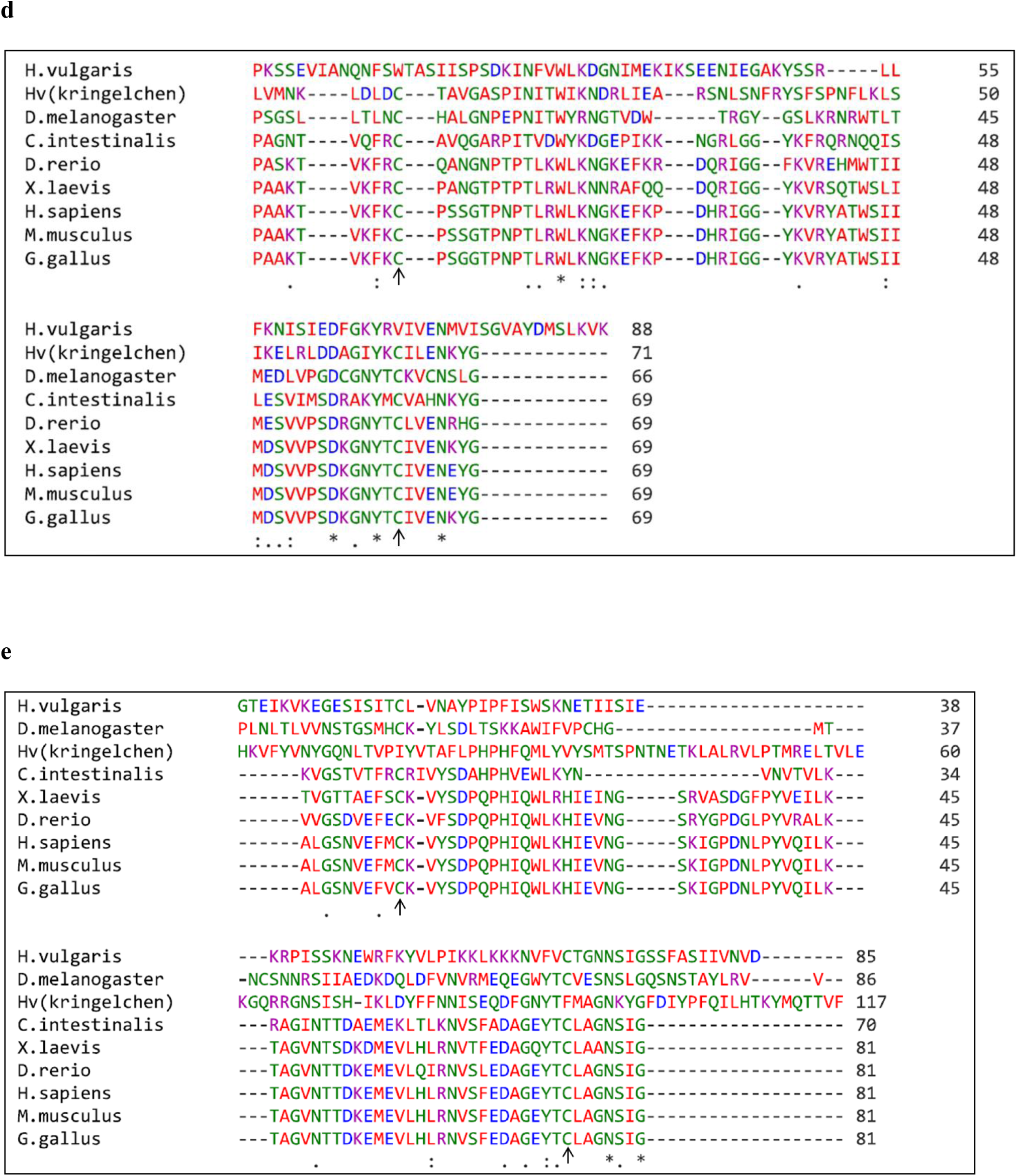
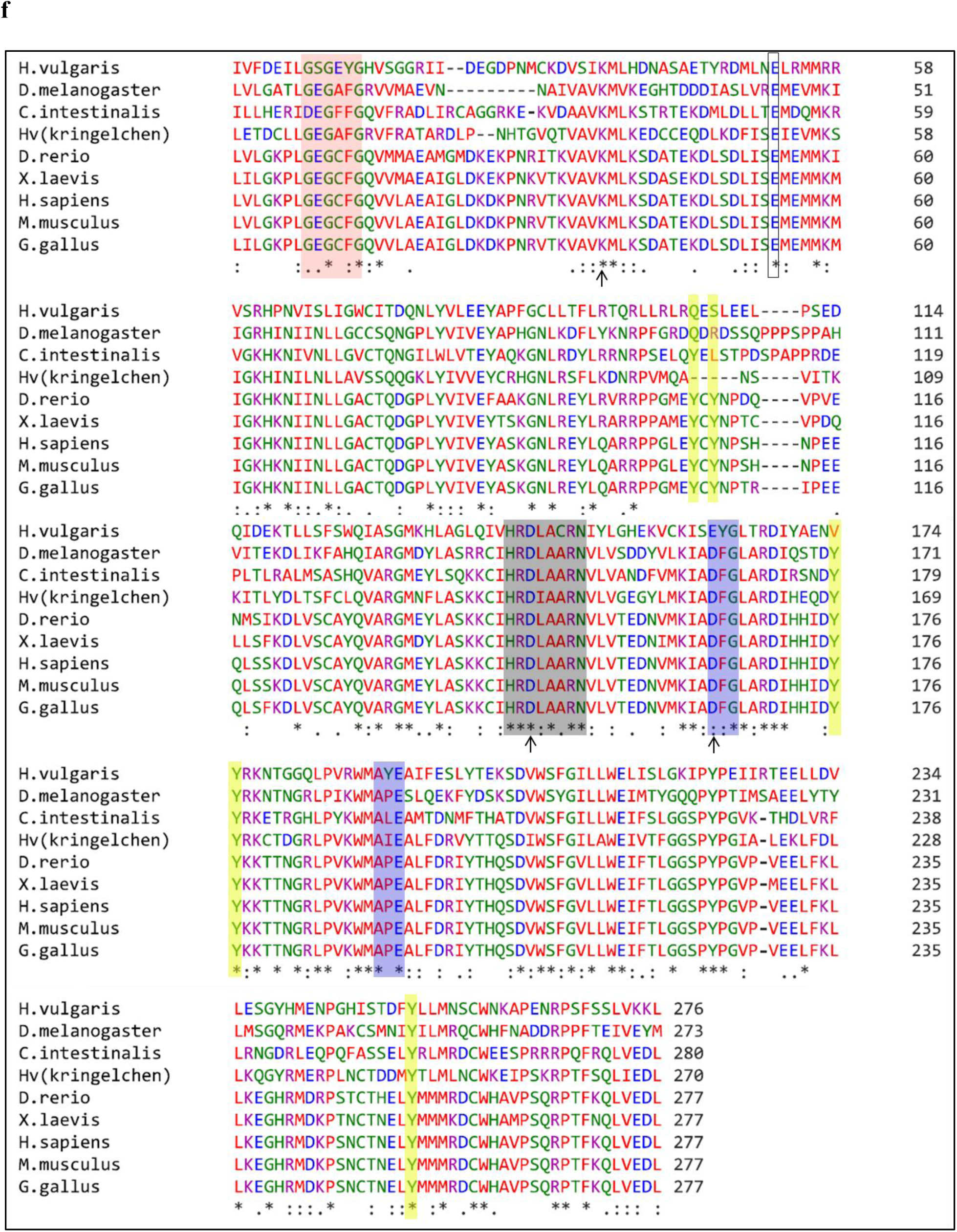
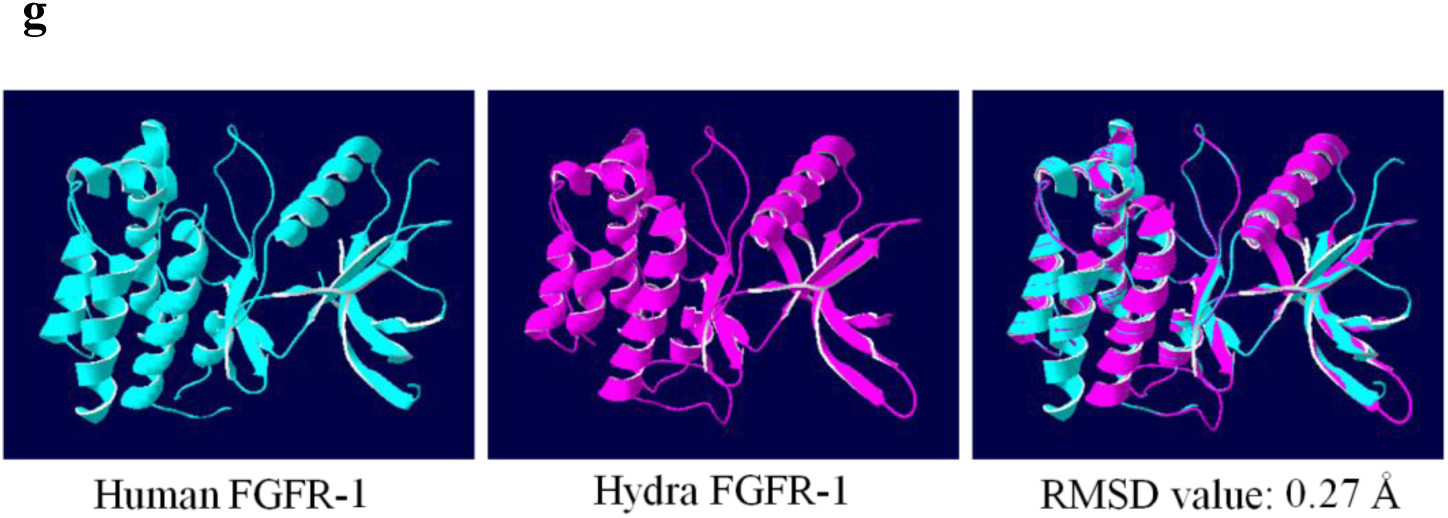
SMART analysis, Tyrosine kinase domain, multiple sequence alignment and homology modelling of FGFR-1. (a) SMART analysis of FGFR-1 protein shows the characteristic Tyrosine kinase domain. (b) The amino acid sequence of the three extracellular immunoglobulin domains, the transmembrane domain and the intracellular tyrosine kinase domain are highlighted in blue, green and red respectively. The region between D1 and D2 highlighted in bold letters denote the serine rich “acid box.” (c), (d), (e) and (f) Multiple sequence alignment of extracellular domains - D1, D2, D3 and tyrosine kinase domain, respectively, with FGFR-1 from other organisms. (*) indicates positions which have a single, fully conserved residue, (:) indicates conservation between groups of strongly similar properties and (.) indicates conservation between groups of weakly similar properties. (c), (d), (e) The conserved cysteines are indicated by arrows. (f) The glycine rich ATP-phosphate binding loop, the catalytic loop, the activation segment residues and tyrosine residues involved in autophosphorylation are highlighted in red, grey, blue and yellow respectively. The box indicates Glutamate residue of the αC helix and the K/D/D motif is indicated with arrows. (g) Tertiary structure of the tyrosine kinase domain of HyFGFR-1 was simulated from available solved structure of Tyrosine kinase domain of human FGFR-1, using Swiss Model Tool. Hydra FGFR-1 and human FGFR-1 were superimposed using Iterative Magic Fit tool in SPDBV (RMSD: 0.27 Å).

Amino acids 50 to 173 of HyFGF-1 form the core FGF domain that consists of 12 antiparallel β strands (β1-β12). These β strands get arranged into three sets of four stranded β sheets and form a folded β trefoil structure [17] (Fig. 2b, d). Unlike other FGF ligands, FGF-1 lacks the N-terminal signal peptide required for the secretion of the protein through the endoplasmic reticulum–golgi secretory pathway (Fig. 2a). Hence, FGF-1 is a paracrine ligand and due to its high affinity for heparan sulphate glycosaminoglycan (HSGAG) acts in a local manner close to its source of secretion [18]. Along with the core FGF domain, the conserved residues also include the three heparan sulphate glycosaminoglycan (HSGAG) binding sites (HBS) of FGF-1. These HBS are rich in basic amino acids – lysine (K) and arginine (R) [19] (Fig. 2c). HSGAGs are known to promote and stabilize ligand-receptor interaction by binding to both FGF-1 and FGFR-1, simultaneously [18]. β10 and β11 loops are a part of the HBS-1, β5, β6 and β9 loops are part of the HBS-2 while β1 loop is a part of HBS-3 (Fig. 2c).

FGFR-1 is a membrane protein with an extracellular domain at the N-terminal, which is made up of three immunoglobulin (IG)-like subdomains – D1, D2 and D3, transmembrane domain made up of a single α helix and an intracellular tyrosine kinase domain [20] (Fig. 4a). Amino acids 31-87 form D1, 110-197 form D2 and 207-291 form D3. Amino acids 300-322 form α helix and amino acids 428 to 703 form the tyrosine kinase domain of HyFGFR-1 (Fig. 4b). Binding of FGF-1 to the extracellular domain of FGFR-1 causes receptor dimerization and trans-autophosphorylation of the intracellular tyrosine kinase domain leading to its activation and downstream signaling. Sequential phosphorylation of six tyrosine residues of FGFR-1 leads to complete activation of the kinase domain. Initially, Y653 is phosphorylated, followed by Y583, Y463, Y766, Y585, Y654 and Y730. Further, phosphorylation of Y677 and Y766 is necessary for STAT 3 and Phospholipase Cγ binding [21]. MSA by clustal omega showed that two of these tyrosine residues – Y-654/Y-730 in vertebrates, correspond to Y-602/Y-679 in hydra and are conserved in HyFGFR-1. Y-583, Y-585 and Y-653 in vertebrate FGFR-1 are replaced by Q-531, S-533 and V-601, respectively, in HyFGFR-1 (Fig. 4c). Other important features of tyrosine kinase domain of vertebrate FGFR-1 are also conserved in HyFGFR-1. These include the glycine rich sequence – GXGXXG (amino acids 435-440) that acts as the ATP-phosphate binding loop, the glutamate residue of the αC helix (E-479), a K/D/D motif (K-462, D-576, D-172), the catalytic loop (amino acids 569-576) and one of the activation segment tyrosines (Y-602). The activation segment in HyFGFR-1 begins with amino acids EYG (amino acids 589-591) (DFG in case of vertebrates) and ends with AYE (amino acids 616-618) (APE in case of vertebrates) (Fig. 4c).

VEGFR-2 is comprised of an extracellular domain, a single transmembrane segment, a juxtamembrane domain and an intracellular tyrosine kinase (TyrKc) domain that contains an insert of 70 amino acids. The extracellular domain is made up of eight immunoglobulin (IG) like domains. The signal peptide is present at the N-terminal (Fig. 6a). Other important residues of VEGFR-2 include a glycine-rich (GXGXXG) ATP-phosphate binding loop, the catalytic loop, the activation segment that begins with amino acids DFG and ends with APE, the activation segment tyrosines, the glutamate residue of the αC helix (E-) and a K/D/D motif. In humans, binding of VEGF to the extracellular domain leads to autophosphorylation of six tyrosine residues. Among these, autophosphorylation of Y-1054 and Y-1059 within the activation loop leads to increased kinase activity. In the active state of VEGFR-2 kinase, K-868 of the K/D/D motif forms ion pairs with the α and β phosphates of ATP and also with E-885 of the αC helix. In the inactive state of the enzyme (in the absence of ATP), K-868 binds to the phosphotyrosine of the activation segment, that is far from E-885. In addition, D-1028 of the catalytic loop positions the tyrosyl group of the substrate protein in a catalytically competent state. D-1046 of the activation loop is also a part of the magnesium binding loop. Thus, D-1046 binds to Mg^2+^ and in turn leads to binding of the α, β and γ phosphate groups of ATP [22]. In case of HyVEGFR-2, amino acids 1-16 form the N-terminal signal peptide. Amino acids 29-115, 129-221, 235-318, 332-429, 445-509, 642-733, 743-851 and 866-931 form the eight IG like domains of the extracellular domain. The transmembrane domain is made up of amino acids 960-982 and amino acids 1032-1349 form the tyrosine kinase domain of HyVEGFR-2 (Fig. 6b). In case of HyVEGFR-2, as observed from MSA, important features like the glycine-rich (GXGXXG) ATP-phosphate binding loop (amino acids 1039-1044), the catalytic loop (amino acids 1216-1223), the activation segment that begins with amino acids DFG (amino acids 1236-1238) and ends with AVE (amino acids 1263-1265), the Glutamate residue (E-1124) of the αC helix and the K/D/D motif are conserved in hydra (K-1107, D-1218, D-1263). In case of the activation segment tyrosines, Y-1054 and Y-1059 are conserved in all phyla studied, except in hydra, where Y-1054 is replaced by H-1244 (Fig. 6c).

### 3.4. Predicted structures of HyFGF-1, HyFGFR-1 and HyVEGFR-2 proteins are similar to their human counterparts

Solved crystal structures of human FGF-1, FGFR-1 and VEGFR-2 were used as templates for simulation of the respective tertiary structures of hydra using Swiss model tool. The structure of human FGF-1, available in PDB (PDB ID: 4q9g.1), has been determined by X-ray diffraction with a resolution of 1.55 Å [23] and was used as a template for model building. The simulated hydra model, when superimposed on human FGF-1 using Iterative Magic Fit tool in SPDBV, gave a RMSD value of 0.35 Å indicating similarity with human FGF-1 (Fig. 2d). HyFGF-1 is composed of 12 β-sheets similar to human FGF-1.

Structure of tyrosine kinase domain of human FGFR-1 determined by X-ray diffraction with a resolution of 2.40 Å [24] has been deposited in PDB (PDB ID: 1agw.1). This structure was used as a template for HyFGFR-1 model building. The simulated hydra model comprises of 8 β-sheets and 8 α-helices while human structure has 8 β-sheets and 9 α-helices. The simulated hydra model was superimposed on human (why is human sometimes capitalized? correct this everywhere) FGFR-1 and resulted in RMSD value of 0.27 Å, indicating close similarity between tyrosine kinase domain of hydra and human FGFR-1 (Fig. 4d).

Tyrosine kinase domain structure of human VEGFR-2 has been determined using X-ray diffraction with a resolution of 1.64 Å [25]. This structure, available in PDB (PDB ID: 3vnt.1), was used as a template to build the model for HyVEGFR-2. Superimposition of the simulated HyVEGFR-2 tyrosine kinase domain on human VEGFR-2 resulted in RMSD value of 0.37 Å indicating similarity between the two proteins. The simulated tyrosine kinase domain of hydra model is made up of 10 β-sheets and 10 α-helices as opposed to 10 β-sheets and 9 α-helices in human VEGFR-2 (Fig. 6d).

### 3.5. HyFGF-1, HyFGFR-1 and HyVEGFR-2 cluster with invertebrates in phylogenetic analysis

Annotated protein sequences of FGF-1, FGFR-1 and VEGFR-2 from different organisms were retrieved from Uniprot and NCBI databases. These sequences were compared with HyFGF-1, HyFGFR-1 and HyVEGFR-2 using NCBI BLASTp tool to determine the homologous sequences. Among the sequences compared, HyFGF-1 showed highest identity of 37% and similarity of 52% with Chick FGF-1 and 29% Identity and 49% similarity to *Nematostella* FGF-1. HyFGFR-1 showed 40% identity and 63% similarity with FGFR-1 from vertebrates and 39% identity and 58% similarity with FGFR-1 from invertebrates. HyVEGFR-2 exhibited 50% identity and 70% similarity with *Podocoryna* VEGFR-2 and 39% identity and 52 % similarity with vertebrate VEGFR-2.

Phylogenetic analysis was conducted in MEGA 7.0 software. FGF-1, FGFR-1 and VEGFR-2 sequences from different organisms were aligned using MUSCLE program. The evolutionary history was inferred using the Neighbor-Joining method and the bootstrap test was set to 5000 replicates. The evolutionary distances were computed using the Poisson correction method and are in the units of the number of amino acid substitutions per site. All positions containing gaps and missing data were eliminated. There were a total of 101, 253 and 248 positions in the final dataset of FGF-1, FGFR-1 and VEGFR-2 respectively. The analysis showed that HyFGF-1, HyFGFR-1 and HyVEGFR-2 clustered together with their invertebrate counterparts followed by vertebrate sequences. The respective protein sequences of *Amphimedon queenslandica*, member of phylum Porifera, were used as an outgroup (Figs. 3, 5, 7).

**Fig. 5.**
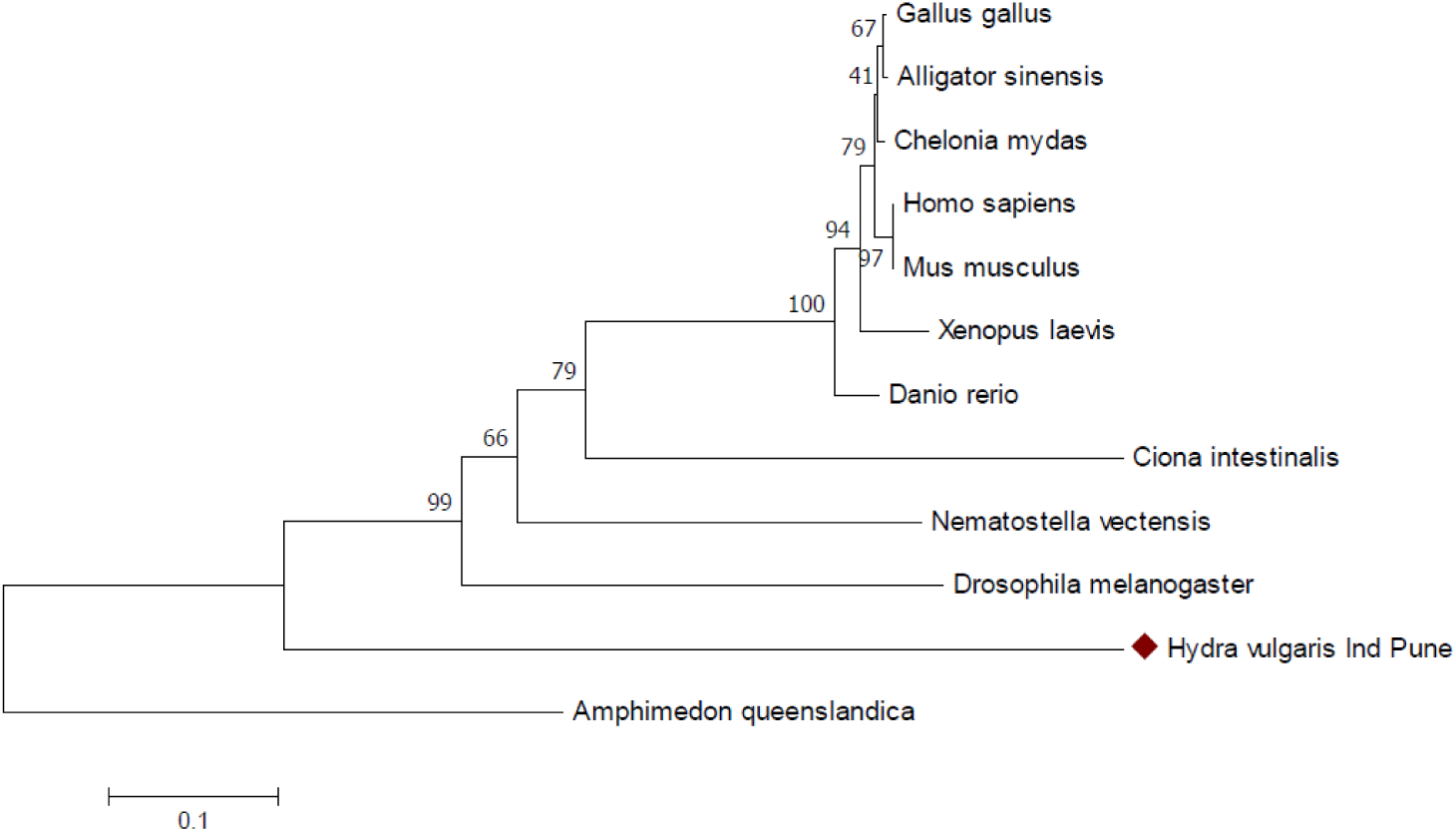
Phylogenetic analysis of FGFR-1 conducted in MEGA7. The evolutionary history was inferred using the Neighbor-Joining method. The optimal tree with the sum of branch length = 2.26881742 is shown. The percentage of replicate trees in which the associated taxa clustered together in the bootstrap test (5000 replicates) are shown next to the branches. The tree is drawn to scale, with branch lengths in the same units as those of the evolutionary distances used to infer the phylogenetic tree. The evolutionary distances were computed using the Poisson correction method and are in the units of the number of amino acid substitutions per site. The analysis involved 12 amino acid sequences. All positions containing gaps and missing data were eliminated. There were a total of 253 positions in the final dataset.

**Fig. 6.**
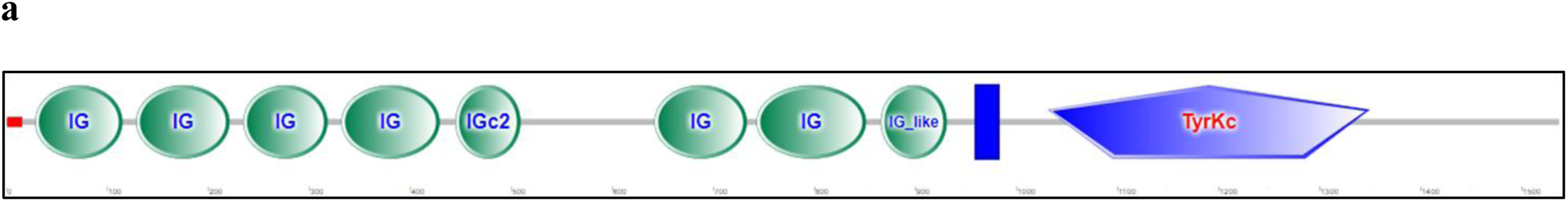
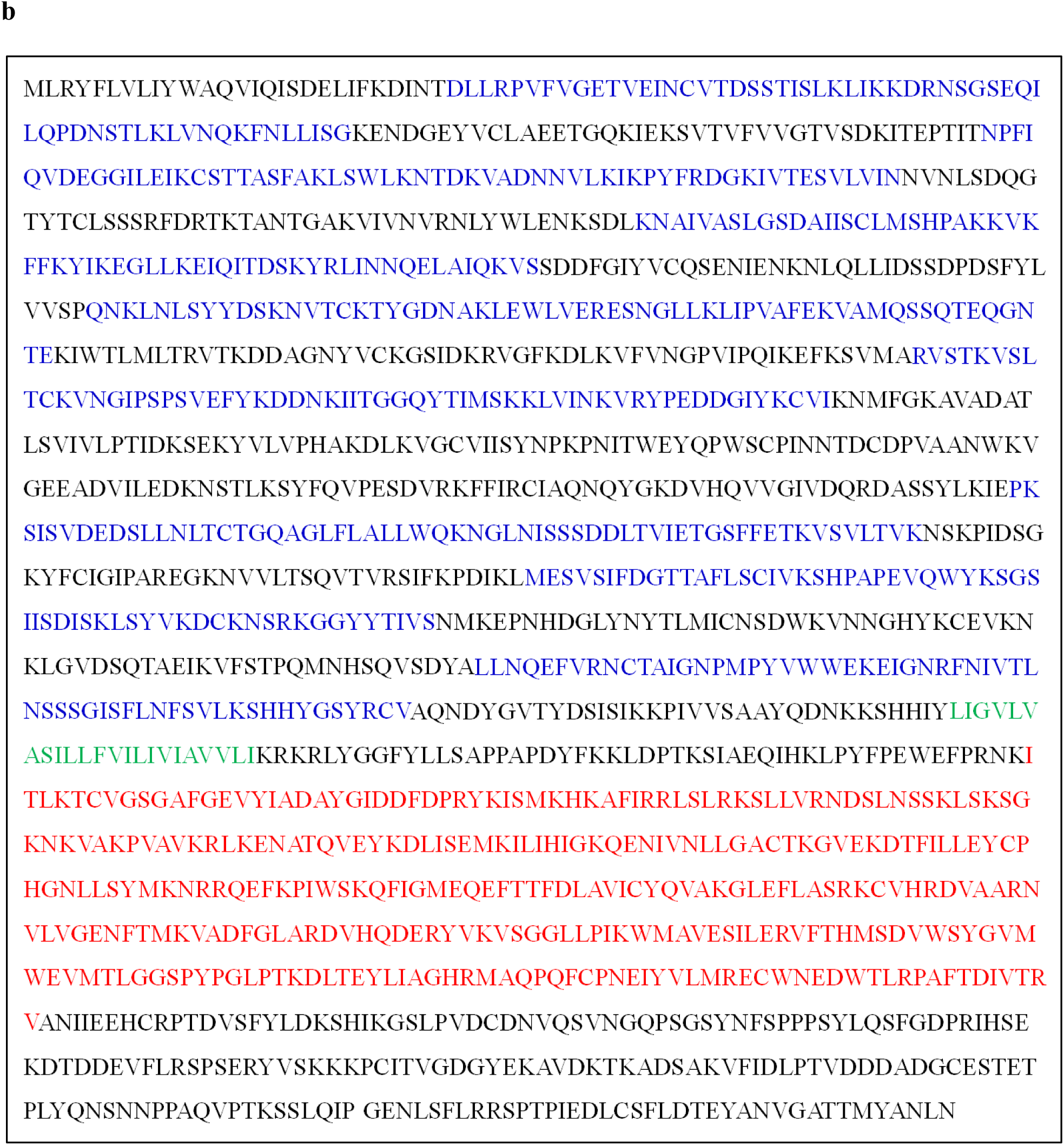
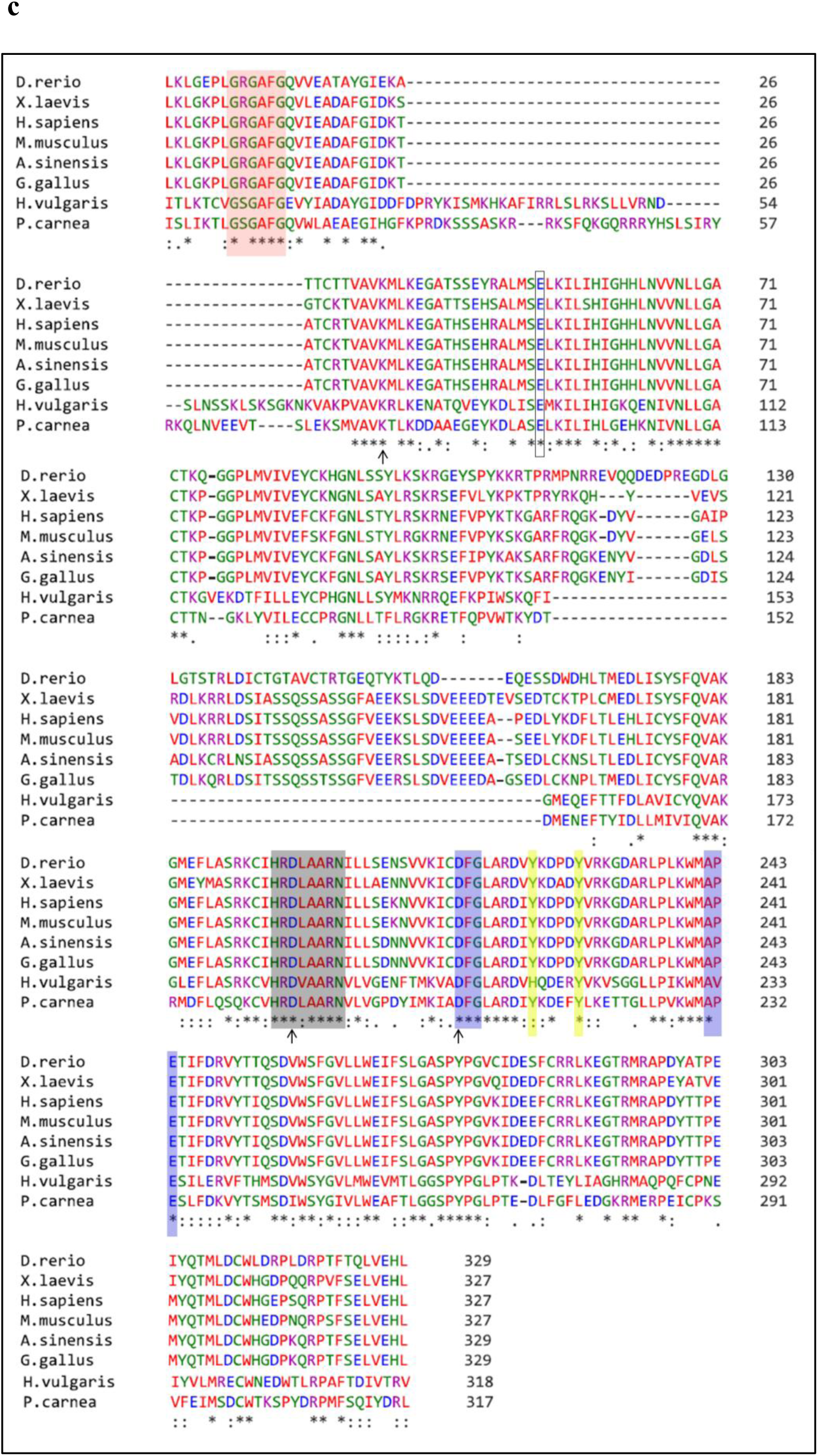
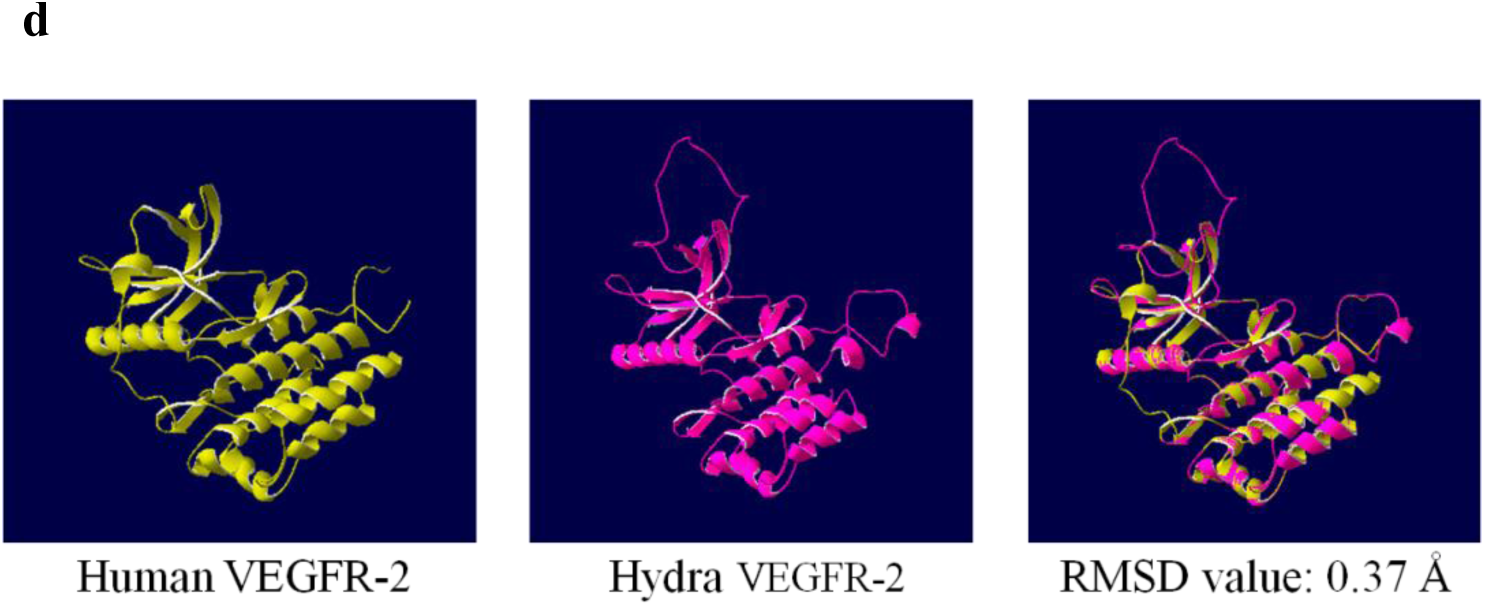
SMART analysis, Tyrosine kinase domain, multiple sequence alignment and homology modelling of VEGFR-2 tyrosine kinase domain. (a) SMART analysis of VEGFR-2 protein shows the characteristic Tyrosine kinase domain. (b) The amino acid sequence of the eight extracellular immunoglobulin domains, the transmembrane domain and the intracellular tyrosine kinase domain are highlighted in blue, green and red respectively. (c) Multiple sequence alignment of tyrosine kinase domains of hydra VEGFR-2 and VEGFR-2 from other organisms shows the conservation of amino acid residues – (*) indicates positions which have a single, fully conserved residue, (:) indicates conservation between groups of strongly similar properties and (.) indicates conservation between groups of weakly similar properties. (d) Tertiary structure of tyrosine kinase domain of HyVEGFR-2 was simulated from available solved structure of the tyrosine kinase domain of human VEGFR-2, using Swiss Model Tool. Hydra VEGFR-2 and human VEGFR-2 were superimposed using Iterative Magic Fit tool in SPDBV (RMSD: 0.37 Å).

**Fig. 7.**
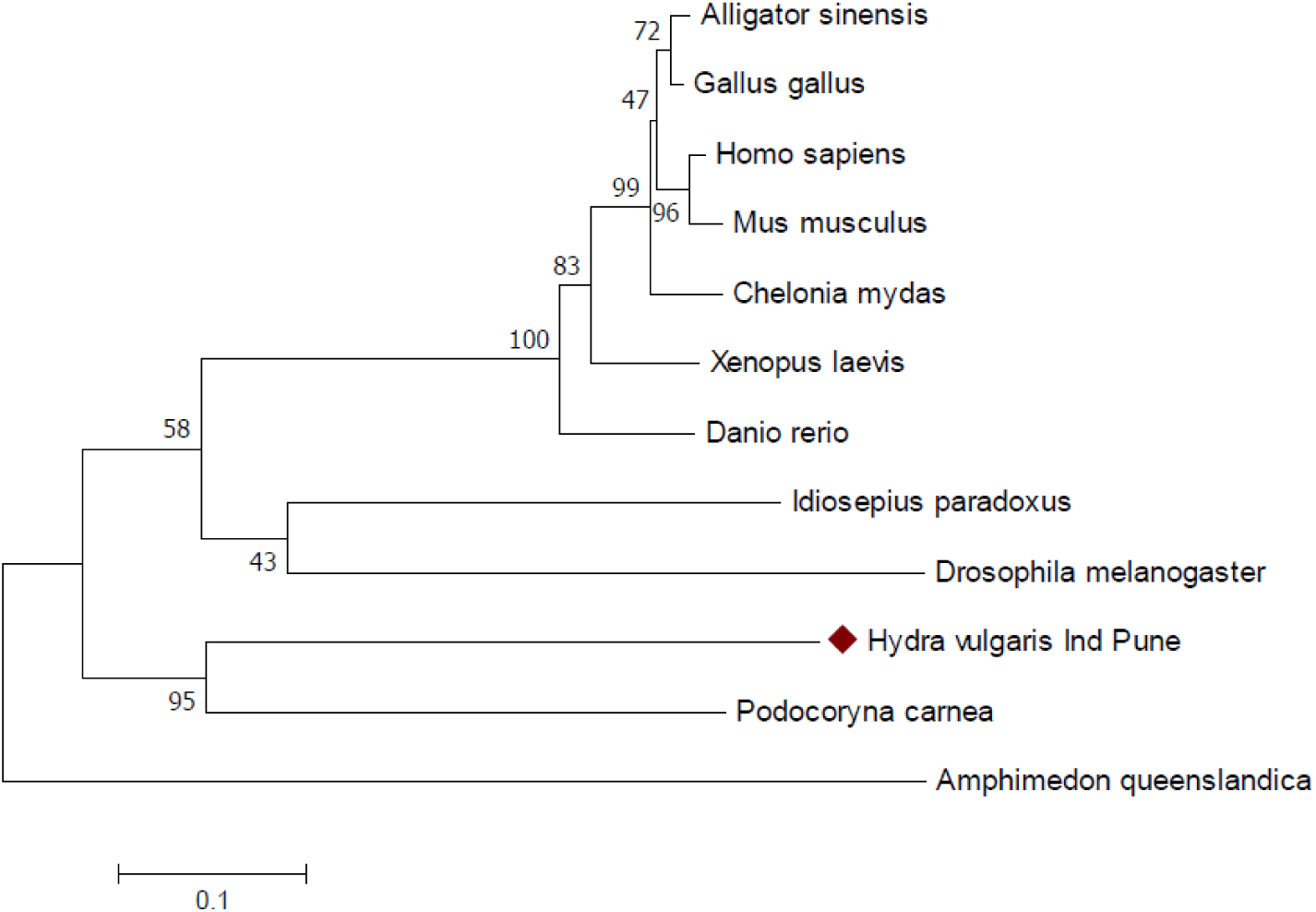
Phylogenetic analysis of VEGFR-2 conducted in MEGA7. The evolutionary history was inferred using the Neighbor-Joining method. The optimal tree with the sum of branch length = 2.38415694 is shown. The percentage of replicate trees in which the associated taxa clustered together in the bootstrap test (5000 replicates) are shown next to the branches. The tree is drawn to scale, with branch lengths in the same units as those of the evolutionary distances used to infer the phylogenetic tree. The evolutionary distances were computed using the Poisson correction method and are in the units of the number of amino acid substitutions per site. The analysis involved 12 amino acid sequences. All positions containing gaps and missing data were eliminated. There were a total of 248 positions in the final dataset

### 3.6. Expression of *FGF-1, FGFR-1* and *VEGFR-2* in hydra

The expression of *FGF-1, FGFR-1* and *VEGFR-2* transcripts in hydra was studied by whole mount *in situ* hybridization. *FGF-1* was localized specifically to the endoderm of the basal disc and tentacles (Fig. 8a, b). *FGFR-1* showed strong expression in the endoderm of the body column and a weak expression in endoderm of the tentacles (Fig. 8c, d). *VEGFR-2* was found to be expressed in endoderm of the tentacles and body column, with the expression decreasing from foot to head (Fig. 8e, f).

**Fig. 8.**
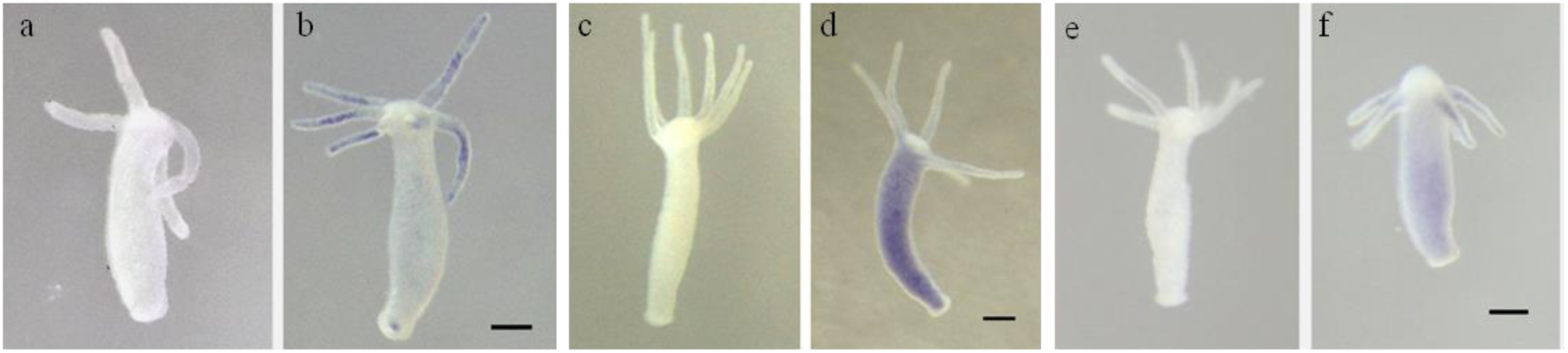
Localization of *FGF-1, FGFR-1* and *VEGFR-2* in hydra by whole mount *in situ* hybridization. Whole mount *in situ* hybridization with DIG labeled FGF-1 antisense riboprobes (b) shows expression in the endoderm of the basal disc and tentacles. FGFR-1 transcripts are strongly expressed in the endoderm of body column with weak expression in endoderm of the tentacles. (d). VEGFR-2 transcripts are localized in endoderm of the tentacles and body column, with the expression decreasing from foot to head (f). (a, c and e) Hybridization with corresponding sense probes for *FGF-1, FGFR-1* and *VEGFR-2*. Scale bar, 200 µm.

### 3.7. Delay in head regeneration upon treatment with VEGFR inhibitor (SU5416) and FGFR inhibitor (SU5402)

In order to examine if VEGF and FGF signaling is involved in hydra head regeneration, the effects of SU5416 and SU5402 on regenerating hydra were studied. Hydra were decapitated and allowed to regenerate the head for 48 hrs in presence of either SU5416 or SU5402. The inhibitor solution was replaced with fresh inhibitor solution after 24 hrs. Hydra kept in hydra medium served as master controls while those in solution with appropriate DMSO concentration served as solvent controls. After treatment for 48 hrs, the polyps were transferred to fresh hydra medium for recovery and left in hydra medium for a further 48 hrs. Head regeneration was found to be completely inhibited after the initial 48 hr-treatment. Post-recovery for 48 hrs, about 70% of the SU5402-treated polyps were able to regenerate a head, whereas only 10% of SU5416-treated polyps were able to regenerate a head (Fig. 9). Thus, treatment with the inhibitors resulted in delayed head regeneration, the effects were partially reversible to different extents, and inhibitor of VEGFR resulted in more potent inhibition of head regeneration.

**Fig. 9.**
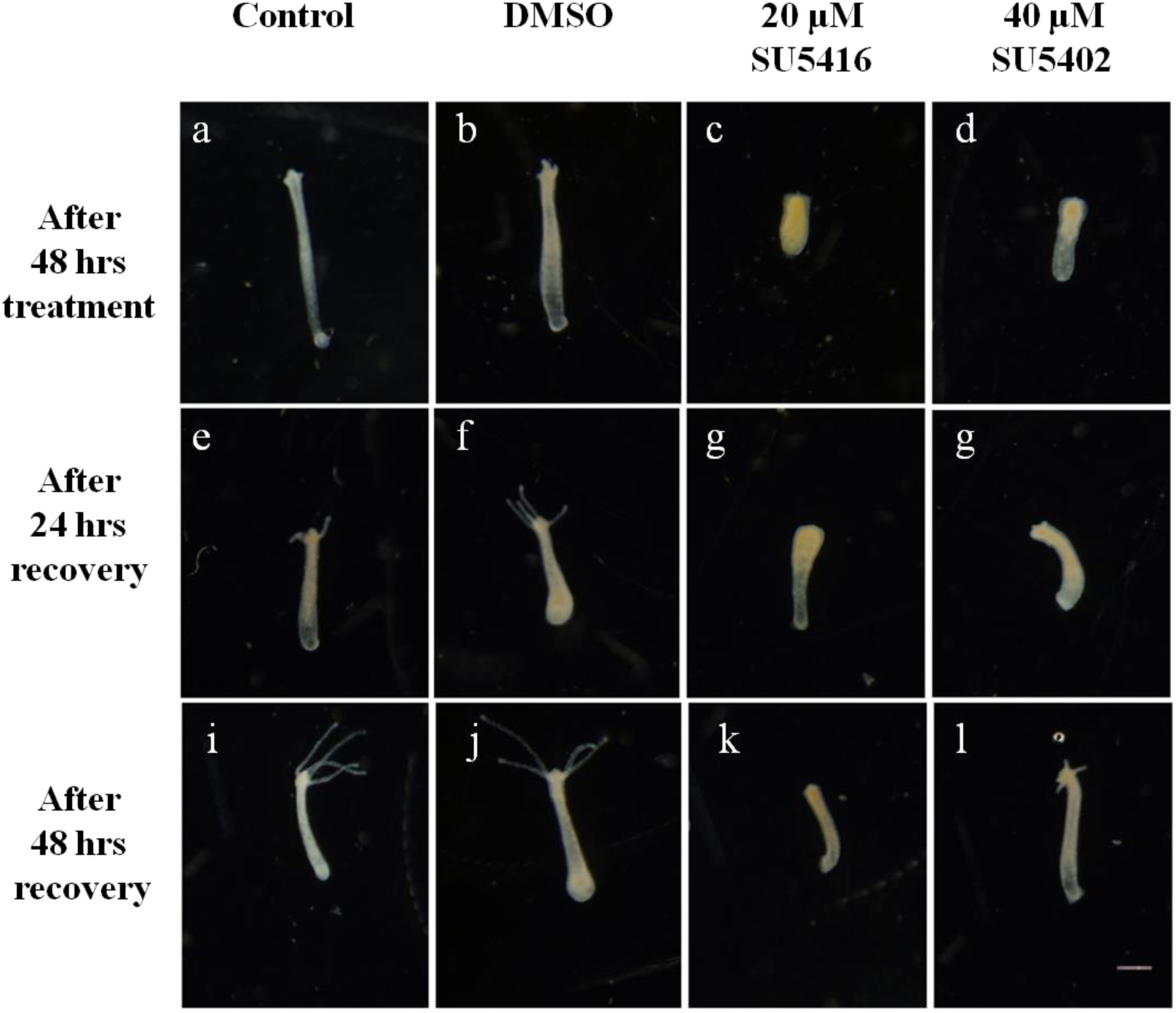
Delay in head regeneration due to SU5416 and SU5402 treatment. Upon 48 hrs treatment of decapitated hydra with VEGFR inhibitor (SU5416) (c) and FGFR inhibitor (SU5402) (d), head regeneration was inhibited. Medium control (a) and solvent control (b) hydra polyps showed normal head regeneration. After 24 hrs and 48 hrs recovery in fresh hydra medium, treated hydra showed signs of delayed head regeneration (g, h, k, l). Polyps used as medium control and solvent control showed complete head regeneration after 48 hrs in fresh hydra medium (e, f, i, j). Scale bar, 200 µm.

### 3.8. Expression of head and tentacle marker genes in hydra

To monitor the process of head regeneration after SU5402 and SU5416 treatment at molecular level, the expression of head specific genes *HyBra1*and *HyKs1* and tentacle specific gene *HyAlx* were studied. *HyAlx* was expressed at the base of the tentacles (Fig. 10a, b), *HyKs1* transcripts were localized in the tentacle zone and at the base of the tentacles (Fig. 10c, d) and *HyBra1* was expressed in the head region (Fig. 10e, f).

**Fig. 10.**
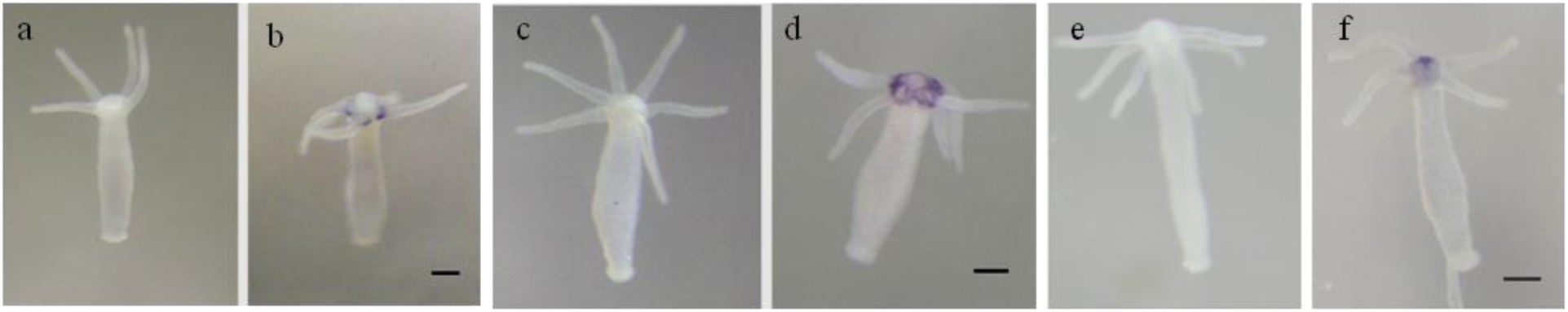
Localization of tentacle specific and head specific genes in hydra by whole mount *in situ* hybridization. Whole mount *in situ* hybridization with DIG labeled *HyAlx* antisense riboprobes (b) shows expression at the base of the tentacles. *HyKs1* transcripts are expressed in the tentacle zone and at base of the tentacles (d). *HyBra1* transcripts are localized in the head region (f). (a, c and e) Hybridization with corresponding sense probes for *HyAlx, HyKs1* and *HyBra1*. Scale bar, 200 µm.

### 3.9. Expression of *HyAlx, HyBra1* and *HyKs1* post SU5402 and SU5416 treatment indicate delay in head regeneration

Decapitated hydra were treated with SU5402 and SU5416 for 48 hrs and the expression of *HyBra1, HyKs1* and *HyAlx* was studied. Expression of these markers indicates the extent of regeneration of head and tentacles. In master and DMSO controls, the marker genes were expressed at optimum levels, whereas in the case of inhibitor treated polyps, the expression of the marker genes was significantly reduced. The expression patterns of these marker genes in treated hydra confirm a delay in head regeneration (Fig. 11).

**Fig. 11.**
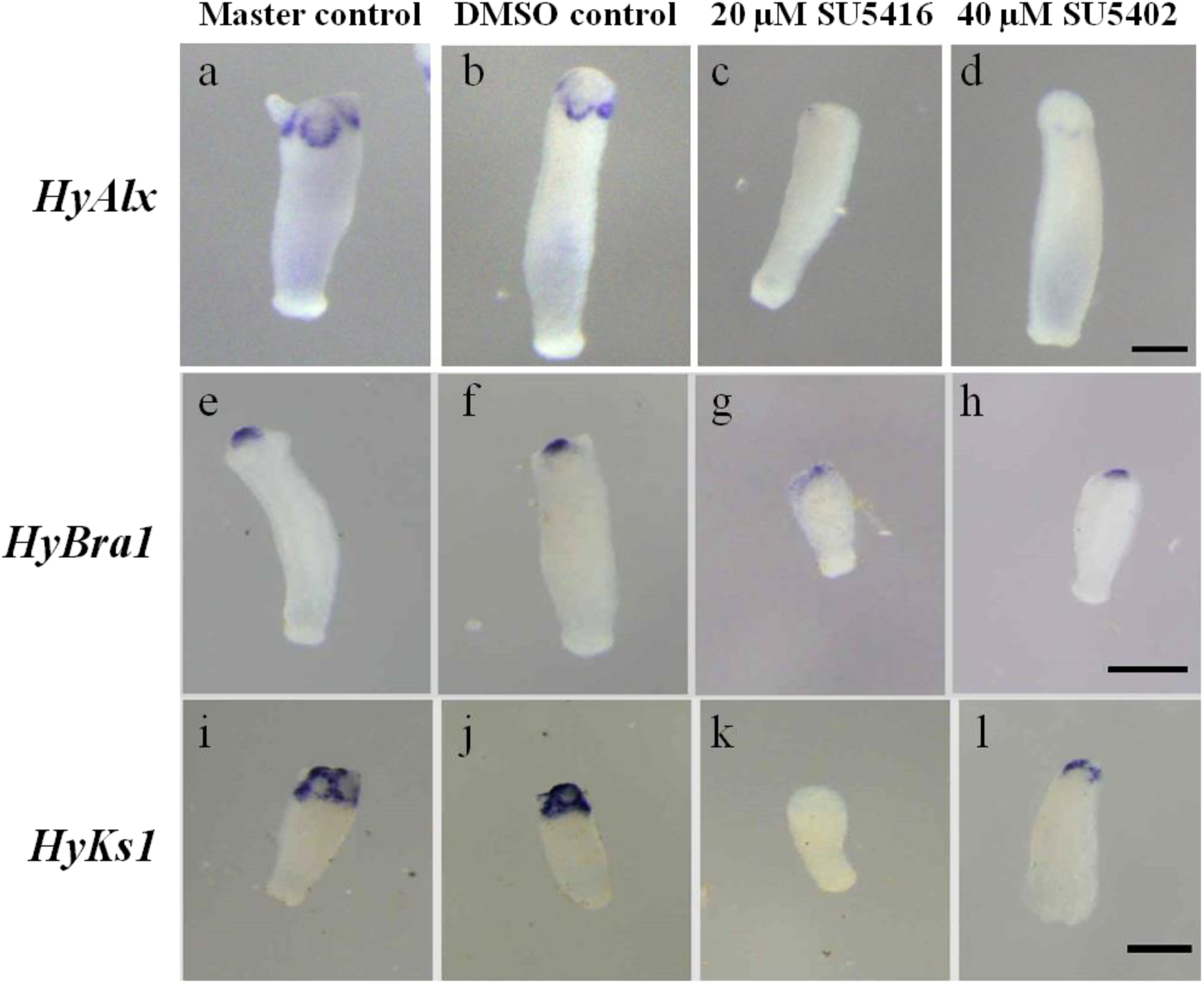
Effect of SU5416 and SU5402 treatment on head regeneration. After 48 hrs of treatment with VEGFR (SU5416) and FGFR (SU5402) inhibitors, the expression of tentacle specific marker HyAlx and head specific markers – HyBra1 and HyKs1 were observed using whole mount *in situ* hybridization. Expression of these markers indicates a head specific and tentacle specific signal. Hence, the expression pattern of HyAlx, HyBra1 and HyKs1 observed in treated hydra (c, d, g, h, k, l) as compared to control hydra (a, b, e, f, i, j), suggests a delay in head regeneration. Scale bar, 200 µm

## 4. Discussion

Hydra offers a unique model system to study head regeneration due to its cellular dynamics. Regeneration of head following decapitation is termed as apical head regeneration. The apical end of hydra consists of mainly the proliferating progenitors that give rise to the differentiated cells. Injury in this region leads to remodeling of the existing tissue to regenerate the lost head [2]. After decapitation, the ectoderm and endoderm stretch over the open wound and form the round shaped apical cap. Tentacle buds appear in the tentacle zone within 30-36 hpd. Further, the apical cap forms a dome shaped structure from which arises the hypostome [8]. To begin with, morphological changes during the process of apical head regeneration were monitored (Fig. 1).

Understanding the role of angiogenic molecules, such as, FGF-1, FGFR-1 and VEGFR-2 in hydra is of particular interest since this can decipher their functions in diploblastic animals. Fibroblast growth factors (FGFs) belong to a family of extracellular secreted proteins and are expressed in nearly all tissues. They play a role during embryonic as well as adult life and are involved in cell proliferation, angiogenesis, organogenesis, tissue repair and regeneration, metabolism, etc. [21]. Previous studies on FGFs in Cnidarians have predicted FGFs in the starlet sea anemone *Nematostella vectensis* and the branching stony coral *Acropora millepora*. Upon phylogenetic analysis, four of the *Nematostella* FGFs grouped with FGF8/17/18/24 subfamily and six grouped with the F1/2 subfamily but with low support. Also one among the four FGF genes, predicted in *Hydra magnipapillata*, belongs to FGF8/17/18/24 subfamily. The rest of the FGFs might belong to the FGF 1/2 subfamily [26, 27]. Partial CDS of FGF-2 from *Hydra vulgaris* Ind-Pune has been reported previously [16]. To our knowledge, this is the first report on the presence of FGF-1 in diploblasts. FGF-1, also known as acidic fibroblast growth factor or heparin binding growth factor-1, belongs to FGF-1 subfamily and is paracrine in its action. FGF-1 generally remains bound to HSGAGs that control its secretion by diffusion through the extracellular matrix. HSGAGs also determine the binding affinity and specificity of FGF with FGF receptors (FGFRs) [21]. *In silico* analysis shows that features of vertebrate FGF-1 are conserved in HyFGF-1. In HyFGF-1, the HSGAG binding sites (HBS) form a contiguous, positively charged surface owing to the presence of lysine (K) and arginine (R) residues (Fig. 2C). Lack of signal peptide in HyFGF-1 implies its paracrine nature (Fig. 2A). HyFGF-1 is composed of 12 β-sheets and adopts the β-trefoil structure characteristic of vertebrate FGF-1 with RMSD value of 0.35 Å (Fig. 2D). Therefore, HyFGF-1 shows considerable structural similarities with FGF-1 from vertebrates.

FGFR-1 belongs to the fibroblast growth factor receptor (FGFR) family of tyrosine kinase receptors. In vertebrates, FGFR receptor is made up of three extracellular immunoglobulin domains (D1-D3), a transmembrane domain and an intracellular tyrosine kinase domain. A serine rich linker sequence called the “acid box” is present between D1 and D2. All these features are present in HyFGFR-1 (Fig. 4a, b). The D2-D3 region is responsible for binding of the FGF ligands and the D1 and “acid box” act in receptor auto-inhibition. Moreover, HSGAG is required for FGF signaling because it leads to FGF-FGFR dimerization by binding to both FGF and FGFR simultaneously [17]. The binding specificity of FGFs and FGFRs is determined by the N-terminal sequence and length of the β-1 strand of FGFs. Also the D3 domain of FGFRs undergoes splicing and determines the binding specificity [18].

Presence of FGFR like tyrosine kinase *kringelchen* has been reported from *Hydra vulgaris* [14]. *Kringelchen* was shown to be involved in boundary formation and tissue constriction which is required for bud detachment. *Kringelchen* showed an overall identity of 26% with HyFGFR-1 while the tyrosine kinase domain of both the receptors was found to be 38% identical. On the other hand, the identity between tyrosine kinase domain of HyFGFR-1 and FGFR-1 from vertebrates was found to be ∼41%, indicating marginally higher similarity with vertebrates (Fig. 4f). The extracellular IG like domains D1, D2 and D3 of *kringelchen* exhibited an identity of 25%, 24% and 12% respectively with HyFGFR-1. D1, D2 and D3 of vertebrate FGFRs showed 16%, 28% and 21% identity with HyFGFR-1. Since D2 and D3 determine the ligand specificity, it is likely that the FGFR-1 in hydra may be activated by FGFs in a manner similar to that in vertebrates. As suggested earlier [14], the conserved cysteines in D1, D2 and D3 are responsible for the formation of Ig like loops. In case of HyFGFR-1, these cysteine residues – C-40 and C-80 in D1and C-222 and C-273 in D3 are conserved while in D2, the conserved cysteines are replaced by the W-123 and V-178 (Fig. 4c, d, e). This could mean that even in the absence of cysteine residues, D2 gets folded into Ig like loop due to the substitution by hydrophobic amino acid residues. Conversely, in kringelchen, the D1 and D2 cysteines are conserved while D3 cysteines are replaced by isoleucine and phenylalanine. Important features of the tyrosine kinase domain of vertebrate receptor tyrosine kinases [28] are conserved in HyFGFR-1 (Fig. 4f). Also the structural conservation between HyFGFR-1 and vertebrate FGFR-1 is evident from the low RMSD value of 0.27 Å in homology modelling (Fig. 4g). These include activation segment tyrosines-Y-602 and Y-679 that are required for autophosphorylation and activation of the kinase domain. These tyrosine residues are also conserved in kringelchen. The second tyrosine of the activation segment is replaced by valine – V-601. This indicates that hydra FGFR-1 may engage tyrosine residues at different positions for trans autophosphorylation and activation of the tyrosine kinase domain. The activation segment in vertebrates begins with DFG and ends with APE. In HyFGFR-1 activation segment begins with EYG and shows conserved substitution by replacing the amino acids DF with EY. The hydrophobic amino acid residue proline in APE is replaced by a hydrophobic tyrosine residue (AYE) in HyFGFR-1. Also in case of kringelchen and FGFR-1 from sea squirt *Ciona*, the APE motif is replaced by hydrophobic residues isoleucine (AIE) and leucine (L) ALE, respectively (Fig. 4f). Thus, substitutions in the APE motif may not hamper its activity.

VEGFR-2 belongs to the type V subfamily of receptor tyrosine kinases. VEGF receptor in vertebrates is made up of seven extracellular immunoglobulin like domains, a single transmembrane domain, a juxtamembrane domain and an intracellular protein–tyrosine kinase domain [22]. HyVEGFR-2 exhibits similar features except that it consists of eight extracellular immunoglobulin (Ig) like domains (Fig. 6b). In cnidarians, homologs of VEGF and VEGFR have been previously reported from the jelly fish *Podocoryne carnea* [29]. HyVEGFR-2 shares 52% identity with the tyrosine kinase domain of *Podocoryne* VEGFR. Most of the features of tyrosine kinase domain of vertebrate VEGFRs, important for signal transduction, are conserved in HyVEGFR-2 (Fig. 6c). In HyVEGFR-2, one of the activation segment tyrosines is replaced by histidine, which could indicate that HyVEGFR-2 employs different tyrosine residues for autophosphorylation. Also, the APE motif of activation segment is replaced by AVE in HyVEGFR-2 (Fig. 6c). The substitution of one hydrophobic residue (proline) with another (valine) may not affect the activity of HyVEGFR-2. The Ig like domains D2 and D3 are responsible for the VEGF-A (ligand) binding [30]. Blocking of D2 and D3 domains leads to inhibition of ligand binding, receptor dimerization, and receptor kinase activation of VEGFR-2 [31]. D2 of HyVEGFR-2 exhibited ∼16% identity with D2 of *Podocoryna* and human VEGFR-2, while domain D3 of HyVEGFR-2 shared 27% and 18% identity with *Podocoryna* and human VEGFR-2 D3 domain, respectively (data not shown). Therefore, the mechanism of ligand binding amongst cnidarians may be conserved but may be different than that in vertebrates. The pair of cysteine residues within each of the seven Ig like domains in vertebrate VEGFR-2 that are responsible for formation of the loop structure, are conserved in HyVEGFR-2. Also a pair of cysteines is present in the eighth domain D8 of HyVEGFR-2 (data not shown). Vertebrate VEGFR-2 is made up of seven Ig like domains while *Podocoryna* VEGFR-2 has six Ig like domains. The D4 and D7 domains were found to be indispensable for ligand mediated receptor signaling [31]. The presence of eighth Ig like domain in HyVEGFR-2 may have a similar function in hydra.

Spatiotemporal expression pattern of a gene provides important insights into its possible function(s) in morphogenesis. Therefore the expression patterns of *HyFGF-1, HyFGFR-1* and *HyVEGFR-2* were studied. *HyFGF-1* shows a very specific expression in the endoderm of the basal disc and endoderm of the tentacles. These regions mainly consist of terminally differentiated cells: differentiated nematocytes in the tentacles and differentiated neurons in both head and foot regions [32]. *HyFGFR-1*, on the other hand, showed ubiquitous expression in the body column. This suggests a possible role of FGF signaling in neurogenesis as the nervous system of hydra is present all over the body in the form of a nerve net. *FGF-2* is expressed in the budding zone of hydra [16]. The expression pattern of *HyFGFR-1* suggests that it could interact with hydra FGF-2. Previously reported FGFR (*kringelchen*) is expressed in both the ectoderm and endoderm of the budding region [14]. Thus, *HyFGF-1* expression suggests that it might not interact with *kringelchen* for FGF signaling in hydra. But *HyFGF-1* expression is similar to that of *FGFf* from *Hydra vulgaris AEP*. It was suggested that FGFf may provide directional cues to the differentiated nematocytes and neurons to migrate towards the terminal regions [15]. HyFGF-1 may also play a similar role.

In vertebrates, VEGF signaling is involved in angiogenesis during embryonic development and also in some physiological processes in adults. In addition, VEGF signaling also regulates cell proliferation and migration, vascular permeability, regeneration, tumour progression, etc. [33]. Homologs of VEGF and VEGFR are present in the jelly fish *Podocoryne carnea* and role of VEGF signaling in tube formation has been proposed [29]. Presence of *VEGF* has been reported in hydra has been reported by us [16]. Localization pattern of *HyVEGFR-2* coincides with the reported expression pattern of *VEGF* in hydra [16] which indicates the possible interaction between the ligand and receptor. Consistent with the ancient role of VEGF signaling in tube formation, *HyVEGFR-2* is expressed in the endoderm of the tentacles (Fig. 8f). Thus, both the components of VEGF signaling VEGF and VEGFR are present in hydra.

To evaluate the role of FGF and VEGF signaling in head regeneration, decapitated hydra were treated with their respective pharmacological inhibitors – SU5402 and SU5416. SU5402 belongs to the class of indolinones that are being used as inhibitors of the tyrosine kinases for cancer therapy. SU5402 acts by competing with ATP for binding to the catalytic domain and inhibits the tyrosine kinase activity of FGFR-1 [34]. SU5416 is a small, lipophilic, highly protein-bound pharmacological inhibitor that functions in a similar way to SU5402 by inhibiting the tyrosine kinase activity of VEGFR [35]. Delayed basal head regeneration and elongation of budding in hydra upon SU5416 treatment has been reported [16]. We find that upon inhibition of the tyrosine kinase domain of FGFR-1 and VEGFR-2 by SU5402 and SU5416, respectively, head regeneration was delayed (Fig. 9). The delay in head regeneration was confirmed by studying the expression of the head and tentacle specific marker genes (Fig. 11). Thus, FGF and VEGF signaling seem to play a role in apical head regeneration.

In conclusion, the present study reports presence of additional components of VEGF and FGF signaling pathways in *Hydra vulgaris* Ind-Pune. The ancient role of FGFs in neurogenesis and VEGFs in tube formation seem to be present in hydra, which represents the basal phylum Cnidaria. The present study strongly suggests roles of VEGF and FGF in tissue regeneration. Since many of the signaling pathways and pattern forming mechanisms are conserved through evolution, similar molecules are likely to participate in tissue and organ regeneration in structurally more complex organisms.

## Abbreviations

VEGF: vascular endothelial growth factor
FGF: fibroblast growth factor
FGFR-1: FGF receptor 1 and VEGFR-2, VEGF receptor 2
A: adenine
T: thymine
G: guanine
C: cytosine
bp: base pairs
cDNA: DNA complementary to RNA
DNA: deoxyribonucleic acid
RNA: ribonucleic acid
DMSO: dimethyl sulfoxide
NCBI: National Center for Biotechnology Information
BLAST: Basic Local Alignment Search Tool
MEGA: Molecular Evolutionary Genetic Analysis
PDB: Protein Data Bank
UniProt: Universal Protein Resource.

## 5. Acknowledgements

This work was supported by intramural funds from MACS-Agharkar Research Institute, Pune, India and Emeritus Scientist Scheme from Council for Scientific and Industrial research (CSIR), New Delhi, India to SG [Grant no. 21(0989)/15/EMR-II]. AT was a recipient of Junior and Senior Research Fellowships (NET) from University Grants Commission, New Delhi [Grant no. 18-12/2011(ii)EU-V]. We are grateful to Dr. Saroj Ghaskadbi, Dr. K.L. Surekha, Dr. Vidya Patwardhan and other lab members for their help and advice during the work.

## References

[1] Agata K, Saito Y, Nakajima E. 2007. Unifying principles of regeneration I: epimorphosis versus morphallaxis. Dev Growth Differ 49: 73–78.

[2] Galliot B and Chera S. 2010. The Hydra model: disclosing an apoptosis-driven generator of Wnt-based regeneration. Trends in Cell Biology 20: 514–523.

[3] Dinsmore C E (Ed.). 1991. A History of Regeneration Research: Milestones in the Evolution of a Science. Cambridge University Press.

[4] Li Q, Yang H, Zhong TP. 2015. Regeneration across metazoan phylogeny: lessons from model organisms. Journal of Genetics and Genomics 42: 57–70.

[5] Bely A E, Nyberg K G. 2010. Evolution of animal regeneration: reemergence of a field. Trends Ecol Evol 25:161–170.

[6] Holstein TW, Hobmayer E, Technau U. 2003. Cnidarians: an evolutionarily conserved model system for regeneration. Developmental Dynamics 226:257-267.

[7] Martínez DE, Bridge D. 2012. Hydra, the everlasting embryo, confronts aging. Int J Dev Biol 56: 479–487.

[8] Bode H R. 2003. Head regeneration in Hydra. Dev Dyn 226: 225–236.

[9] Bosch T C G. 2007. Why polyps regenerate and we don’t: Towards a cellular and molecular framework for Hydra regeneration. Dev Biol 303(2): 421–33.

[10] Schlessinger J. 2000. Cell signaling by receptor tyrosine kinases. Cell 103 (2): 211–225.

[11] Reddy P C, Barve A, Ghaskadbi S. 2011. Description and phylogenetic characterization of common Hydra from India. Curr. Sci. 101: 736–738.

[12] Maddaluno L, Urwyler C, Werner S. 2017. Fibroblast growth factors: key players in regeneration and tissue repair. Development 144: 4047–4060.

[13] Matsumoto K and Ema M. 2014. Roles of VEGF-A signaling in development, regeneration and tumours. J Biochem 156 (1): 1–10.

[14] Sudhop S, Coulier F, Bieller A, Vogt A, Hotz T, Hassel M. 2004. Signaling by the FGFR-like tyrosine kinase, Kringelchen, is essential for bud detachment in *Hydra vulgaris*. Development 131: 4001–4011.

[15] Lange E, Bertrand S, Holz O, Rebscher N, Hassel M. 2014. Dynamic expression of a Hydra FGF at boundaries and termini. Dev Genes Evol 224: 235.

[16] Krishnapati LS, Ghaskadbi S. 2013. Identification and characterization of VEGF and FGF from Hydra. Int J Dev Biol 57: 897–906.

[17] MohammadI M, Olsen SK, Ibrahimi OA. 2005. Structural basis for fibroblast growth factor receptor activation. Cytokine Growth Factor Rev 16(2):107–137.

[18] Beenken A, Mohammadi M. 2009. The FGF family: biology, pathophysiology and therapy. Nat Rev Drug Discov 8(3): 235–253.

[19] Xu R, Ori A, Rudd TR, Uniewicz KA, Ahmed YA, Guimond SE, Skidmore MA, Siligardi G, Yates EA and Fernig DG. 2012. Diversification of the structural determinants of fibroblast growth factor-heparin interactions J Biol Chem 287: 40061– 40073.

[20] Sarabipour S and Hristova K. 2016. Mechanism of FGF receptor dimerization and activation. Nat Commun 7:10262.

[21] Ornitz DM, Itoh N. 2015. The Fibroblast Growth Factor signaling pathway. WIREs Dev Biol 4: 215–266.

[22] Roskoski R Jr. 2008. VEGF receptor protein–tyrosine kinases: Structure and regulation. Biochemical and Biophysical Research Communications 375: 287–291.

[23] Xia X, Longo LM, Blaber M. 2015. Mutation choice to eliminate buried free cysteines in protein therapeutics. J Pharm Sci 104(2): 566–576.

[24] Mohammadi M, McMahon G, Sun L, Tang C, Hirth P, Yeh BK, Hubbard SR, Schlessinger J. 1997. Structures of the tyrosine kinase domain of fibroblast growth factor receptor in complex with inhibitors. Science 276 (5314):955–60.

[25] Okaniwa M, Hirose M, Imada T, Ohashi T, Hayashi Y, Miyazaki T, Arita T, Yabuki M, Kakoi K, Kato J, Takagi T, Kawamoto T, Yao S, Sumita A, Tsutsumi S, Tottori T, Oki H, Sang BC, Yano J, Aertgeerts K, Yoshida S, Ishikawa T. 2012. Design and synthesis of novel DFG-out RAF/vascular endothelial growth factor receptor 2 (VEGFR2) inhibitors. 1. Exploration of [5, 6]-fused bicyclic scaffolds. J Med Chem 55(7): 3452–3478.

[26] Technau U, Rudd S, Maxwell P, Gordon PM, Saina M, Grasso LC, Hayward DC, Sensen CW, Saint R, Holstein TW, Ball EE, Miller DJ. 2005. Maintenance of ancestral complexity and non-metazoan genes in two basal cnidarians. Trends in Genetics 21(12): 633–639.

[27] Matus DQ, Thomsen GH, and Martindale MQ. 2007. FGF signaling in gastrulation and neural development in Nematostella vectensis, an anthozoan cnidarians. Development Genes and Evolution 217: 137–148.

[28] Roskoski R Jr. 2015. A historical overview of protein kinases and their targeted small molecule inhibitors. Pharmacological Research 100: 1–23.

[29] Seipel K, Eberhardt M, Müller P, Pescia E, Yanze N, Schmid V. 2004. Homologs of vascular endothelial growth factor and receptor, VEGF and VEGFR, in the jellyfish Podocoryne carnea. Dev Dyn 231(2):303–12.

[30] Leppänen VM, Prota AE, Jeltsch M, Anisimov A, Kalkkinen N, Strandin T, Lankinen H, Goldman A, Ballmer-Hofer K and Alitalo K. 2010. Structural determinants of growth factor binding and specificity by VEGF receptor 2. PNAS 107(6): 2425–2430.

[31] Hyde CAC, Giese A, Stuttfeld E, Saliba JA, Villemagne D, Schleier T, Binz HK, Hofer KB. 2012. Targeting Extracellular Domains D4 and D7 of Vascular Endothelial Growth Factor Receptor 2 Reveals Allosteric Receptor Regulatory Sites. Molecular and Cellular Biology 32:19: 3802–3813.

[32] Galliot B, Quiquand M, Ghila L, Rosa R, Licina MM, Chera S. 2009. Origins of neurogenesis, a cnidarian view. Developmental Biology 332: 2–24.

[33] Kipryushina YO, Yakovlev KV, Odintsova NA. 2015. Vascular endothelial growth factors: A comparison between invertebrates and vertebrates. Cytokine & Growth Factor Reviews 26: 687–695.

[34] Saradamba A, Buch PR, Murawala HA, Balakrishnan S. 2013. SU5402, a pharmacological inhibitor of fibroblast growth factor receptor (FGFR), effectively hampers the initiation and progression of fin regeneration in teleost fish. European Journal of Zoological Research 2(4):1–9.

[35] O’farrell AM, Yuen HA, Smolich B, Hannah AL, Louie SG, Hong W, Stopeck AT, Silverman LR, Lancet JE, Karp JE, Albitar M, Cherrington JM, Giles FJ. 2004. Effects of SU5416, a small molecule tyrosine kinase receptor inhibitor, on FLT3 expression and phosphorylation in patients with refractory acute myeloid leukemia. Leukemia Research 28: 679–689.

[36] Letunic I and Bork P. 2018. 20 years of the SMART protein domain annotation resource. Nucleic Acids Research 46 (D1); 493–496.

[37] Sievers F, Wilm A, Dineen D, Gibson TJ, Karplus K, Li W, Lopez R, Mcwilliam H, Remmert M, Söding J, Thompson JD, Higgins DG. 2011. Fast scalable generation of high quality multiple sequence alignments using Clustal Omega. Mol. Syst. Biol. 7: 539.

[38] Biasini M, Bienert S, Waterhouse A, Arnold K, Studer G, Schmidt T, Kiefer F, Cassarino T G, Bertoni M, Bordoli L, Schwede T. 2014. SWISS MODEL: modelling protein tertiary and quaternary structure using evolutionary information. Nucleic Acids Res 42: W252–W258.

[39] Guex N and Pietsch MC. 1997. SWISS MODEL and the Swiss-PdbViewer: An environment for comparative protein modelling. Electrophoresis 18: 2714–2723.

[40] Kumar S, Stecher G, Li M, Knyaz C, Tamura K. 2018. MEGA 7: Molecular Evolutionary Genetics Analysis across computing platforms. Molecular Biology and Evolution 35:1547–1549.

